# An improved pig reference genome sequence to enable pig genetics and genomics research

**DOI:** 10.1101/668921

**Authors:** Amanda Warr, Nabeel Affara, Bronwen Aken, H. Beiki, Derek M. Bickhart, Konstantinos Billis, William Chow, Lel Eory, Heather A. Finlayson, Paul Flicek, Carlos G. Girón, Darren K. Griffin, Richard Hall, Greg Hannum, Thibaut Hourlier, Kerstin Howe, David A. Hume, Osagie Izuogu, Kristi Kim, Sergey Koren, Haibou Liu, Nancy Manchanda, Fergal J. Martin, Dan J. Nonneman, Rebecca E. O’Connor, Adam M. Phillippy, Gary A. Rohrer, Benjamin D. Rosen, Laurie A. Rund, Carole A. Sargent, Lawrence B. Schook, Steven G. Schroeder, Ariel S. Schwartz, Ben M. Skinner, Richard Talbot, Elizabeth Tseng, Christopher K. Tuggle, Mick Watson, Timothy P. L. Smith, Alan L. Archibald

## Abstract

The domestic pig (*Sus scrofa*) is important both as a food source and as a biomedical model with high anatomical and immunological similarity to humans. The draft reference genome (Sscrofa10.2) of a purebred Duroc female pig established using older clone-based sequencing methods was incomplete and unresolved redundancies, short range order and orientation errors and associated misassembled genes limited its utility. We present two annotated highly contiguous chromosome-level genome assemblies created with more recent long read technologies and a whole genome shotgun strategy, one for the same Duroc female (Sscrofa11.1) and one for an outbred, composite breed male (USMARCv1.0). Both assemblies are of substantially higher (>90-fold) continuity and accuracy than Sscrofa10.2. These highly contiguous assemblies plus annotation of a further 11 short read assemblies provide an unprecedented view of the genetic make-up of this important agricultural and biomedical model species. We propose that the improved Duroc assembly (Sscrofa11.1) become the reference genome for genomic research in pigs.

## Background

High quality, richly annotated reference genome sequences are key resources and provide important frameworks for the discovery and analysis of genetic variation and for linking genotypes to function. In farmed animal species such as the domestic pig (*Sus scrofa*) genome sequences have been integral to the discovery of molecular genetic variants and the development of single nucleotide polymorphism (SNP) chips [*1*] and enabled efforts to dissect the genetic control of complex traits, including responses to infectious diseases [*2*].

Genome sequences are not only an essential resource for enabling research but also for applications in the life sciences. Genomic selection, in which associations between thousands of SNPs and trait variation as established in a phenotyped training population are used to choose amongst selection candidates for which there are SNP data but no phenotypes, has delivered genomics-enabled genetic improvement in farmed animals [*3*] and plants. From its initial successful application in dairy cattle breeding, genomic selection is now being used in many sectors within animal and plant breeding, including by leading pig breeding companies [*4, 5*].

The domestic pig (*Sus scrofa*) has importance not only as a source of animal protein but also as a biomedical model. The choice of the optimal animal model species for pharmacological or toxicology studies can be informed by knowledge of the genome and gene content of the candidate species including pigs [*6*]. A high quality, richly annotated genome sequence is also essential when using gene editing technologies to engineer improved animal models for research or as sources of cells and tissue for xenotransplantation and potentially for improved productivity [*7, 8*].

The highly continuous pig genome sequences reported here are built upon a quarter of a century of effort by the global pig genetics and genomics research community including the development of recombination and radiation hybrid maps [*9, 10*], cytogenetic and Bacterial Artificial Chromosome (BAC) physical maps [*11, 12*] and a draft reference genome sequence [*13*].

The previously published draft pig reference genome sequence (Sscrofa10.2), developed under the auspices of the Swine Genome Sequencing Consortium (SGSC), has a number of significant deficiencies [*14–17*]. The BAC-by-BAC hierarchical shotgun sequence approach [*18*] using Sanger sequencing technology can yield a high quality genome sequence as demonstrated by the public Human Genome Project. However, with a fraction of the financial resources of the Human Genome Project, the resulting draft pig genome sequence comprised an assembly, in which long-range order and orientation is good, but the order and orientation of sequence contigs within many BAC clones was poorly supported and the sequence redundancy between overlapping sequenced BAC clones was often not resolved. Moreover, about 10% of the pig genome, including some important genes, were not represented (e.g. *CD163*), or incompletely represented (e.g. *IGF2*) in the assembly [*19*]. Whilst the BAC clones represent an invaluable resource for targeted sequence improvement and gap closure as demonstrated for chromosome X (SSCX) [*20*], a clone-by-clone approach to sequence improvement is expensive notwithstanding the reduced cost of sequencing with next-generation technologies.

The dramatically reduced cost of whole genome shotgun sequencing using Illumina short read technology has facilitated the sequencing of several hundred pig genomes [*17, 21, 22*]. Whilst a few of these additional pig genomes have been assembled to contig level, most of these genome sequences have simply been aligned to the reference and used as a resource for variant discovery.

The increased capability and reduced cost of third generation long read sequencing technology as delivered by Pacific Biosciences and Oxford Nanopore platforms, have created the opportunity to generate the data from which to build highly contiguous genome sequences as illustrated recently for cattle [*23, 24*]. Here we describe the use of Pacific Biosciences (PacBio) long read technology to establish highly continuous pig genome sequences that provide substantially improved resources for pig genetics and genomics research and applications.

## Results

Two individual pigs were sequenced independently: a) TJ Tabasco (Duroc 2-14) i.e. the sow that was the primary source of DNA for the published draft genome sequence (Sscrofa10.2) [*13*] and b) MARC1423004 which was a Duroc/Landrace/Yorkshire crossbred barrow (i.e. castrated male pig) from the USDA Meat Animal Research Center. The former allowed us to build upon the earlier draft genome sequence, exploit the associated CHORI-242 BAC library resource (https://bacpacresources.org/ http://bacpacresources.org/porcine242.htm) and evaluate the improvements achieved by comparison with Sscrofa10.2. The latter allowed us to assess the relative efficacy of a simpler whole genome shotgun sequencing and Chicago Hi-Rise scaffolding strategy [*25*]. This second assembly also provided data for the Y chromosome, and supported comparison of haplotypes between individuals. In addition, full-length transcript sequences were collected for multiple tissues from the MARC1423004 animal, and used in annotating both genomes.

### Sscrofa11.1 assembly

Approximately sixty-five fold coverage (176 Gb) of the genome of TJ Tabasco (Duroc 2-14) was generated using Pacific Biosciences (PacBio) single-molecule real-time (SMRT) sequencing technology. A total of 213 SMRT cells produced 12,328,735 subreads of average length 14,270 bp and with a read N50 of 19,786 bp (Table S1). Reads were corrected and assembled using Falcon (v.0.4.0) [*26*], achieving a minimum corrected read cutoff of 13 kb that provided 19-fold genome coverage for input resulting in an initial assembly comprising 3,206 contigs with a contig N50 of 14.5 Mb.

The contigs were mapped to the previous draft assembly (Sscrofa10.2) using Nucmer [*27*]. The long range order of the Sscrofa10.2 assembly was based on fingerprint contig (FPC) [*12*] and radiation hybrid physical maps with assignments to chromosomes based on fluorescent *in situ* hybridisation data. This alignment of Sscrofa10.2 and the contigs from the initial Falcon assembly of the PacBio data provided draft scaffolds that were tested for consistency with paired BAC and fosmid end sequences and the radiation hybrid map [*9*]. The draft scaffolds also provided a framework for gap closure using PBJelly [*28*], or finished quality Sanger sequence data generated from CHORI-242 BAC clones from earlier work [*13, 20*].

Remaining gaps between contigs within scaffolds, and between scaffolds predicted to be adjacent on the basis of other available data, were targeted for gap filling with a combination of unplaced contigs and previously sequenced BACs, or by identification and sequencing of BAC clones predicted from their end sequences to span the gaps. The combination of methods filled 2,501 gaps and reduced the number of contigs in the assembly from 3,206 to 705. The assembly, Sscrofa11 (GCA_000003025.5), had a final contig N50 of 48.2 Mb, only 103 gaps in the sequences assigned to chromosomes, and only 583 remaining unplaced contigs (Table 1). Two acrocentric chromosomes (SSC16, SSC18) were each represented by single, unbroken contigs. The SSC18 assembly also includes centromeric and telomeric repeats (Tables S2, S3; Figs. S1, S2), albeit the former probably represent a collapsed version of the true centromere. The reference genome assembly was completed by adding Y chromosome sequences from other sources (GCA_900119615.2) [*20*] because TJ Tabasco (Duroc 2-14) was female. The resulting reference genome sequence was termed Sscrofa11.1 and deposited in the public sequence databases (GCA_000003025.6) (Table 1).

**Table 1.** Assembly statistics. Summary statistics for assembled pig genome sequences and comparison with current human reference genome. (source: NCBI, https://www.ncbi.nlm.nih.gov/assembly/; * includes mitochondrial genome.

| Assembly | Sscrofa10.2 | Sscrofa11 | Sscrofa11.1 | USMARCv1.0 | GRCh38.p12 |
| --- | --- | --- | --- | --- | --- |
| Total sequence length | 2,808,525,991 | 2,456,768,445 | 2,501,912,388 | 2,755,438,182 | 3,099,706,404 |
| Total ungapped length | 2,519,152,092 | 2,454,899,091 | 2,472,047,747 | 2,623,130,238 | 2,948,583,725 |
| Number of scaffolds | 9,906 | 626 | 706 | 14,157 | 472 |
| Gaps between scaffolds | 5,323 | 24 | 93 | 0 | 349 |
| Number of unplaced scaffolds | 4,562 | 583 | 583 | 14,136 | 126 |
| Scaffold N50 | 576,008 | 88,231,837 | 88,231,837 | 131,458,098 | 67,794,873 |
| Scaffold L50 | 1,303 | 9 | 9 | 9 | 16 |
| Number of unspanned gaps | 5,323 | 24 | 93 | 0 | 349 |
| Number of spanned gaps | 233,116 | 79 | 413 | 661 | 526 |
| Number of contigs | 243,021 | 705 | 1,118 | 14,818 | 998 |
| Contig N50 | 69,503 | 48,231,277 | 48,231,277 | 6,372,407 | 57,879,411 |
| Contig L50 | 8,632 | 15 | 15 | 104 | 18 |
| Number of chromosomes* | *21 | 19 | *21 | *21 | 24 |

The medium to long range order and orientation of Sscrofa11.1 assembly was assessed by comparison to an existing radiation hybrid (RH) map [*9*]. The comparison strongly supported the overall accuracy of the assembly (Fig. 1a), despite the fact that the RH map was prepared from a cell line of a different individual. There is one major disagreement between the RH map and the assembly on chromosome 3, which will need further investigating. The only other substantial disagreement on chromosome 9, is explained by a gap in the RH map [*9*]. The assignment and orientation of the Sscrofa11.1 scaffolds to chromosomes was confirmed with fluorescent *in situ* hybridisation (FISH) of BAC clones (Table S4, Fig. S3). The Sscrofa11.1 and USMARCv1.0 assemblies were searched using BLAST with sequences derived from the BAC clones which had been used as probes for the FISH analyses. For most BAC clones these sequences were BAC end sequences [*12*], but in some cases these sequences were incomplete or complete BAC clone sequences [*13, 20*]. The links between the genome sequence and the BAC clones used in cytogenetic analyses by fluorescent *in situ* hybridization are summarised in Table S4. The fluorescent *in situ* hybridization results indicate areas where future assemblies might be improved. For example, the Sscrofa11.1 unplaced scaffolds contig 1206 and contig1914 may contain sequences that could be added to end of the long arms of SSC1 and SSC7 respectively.

**Figure 1.**
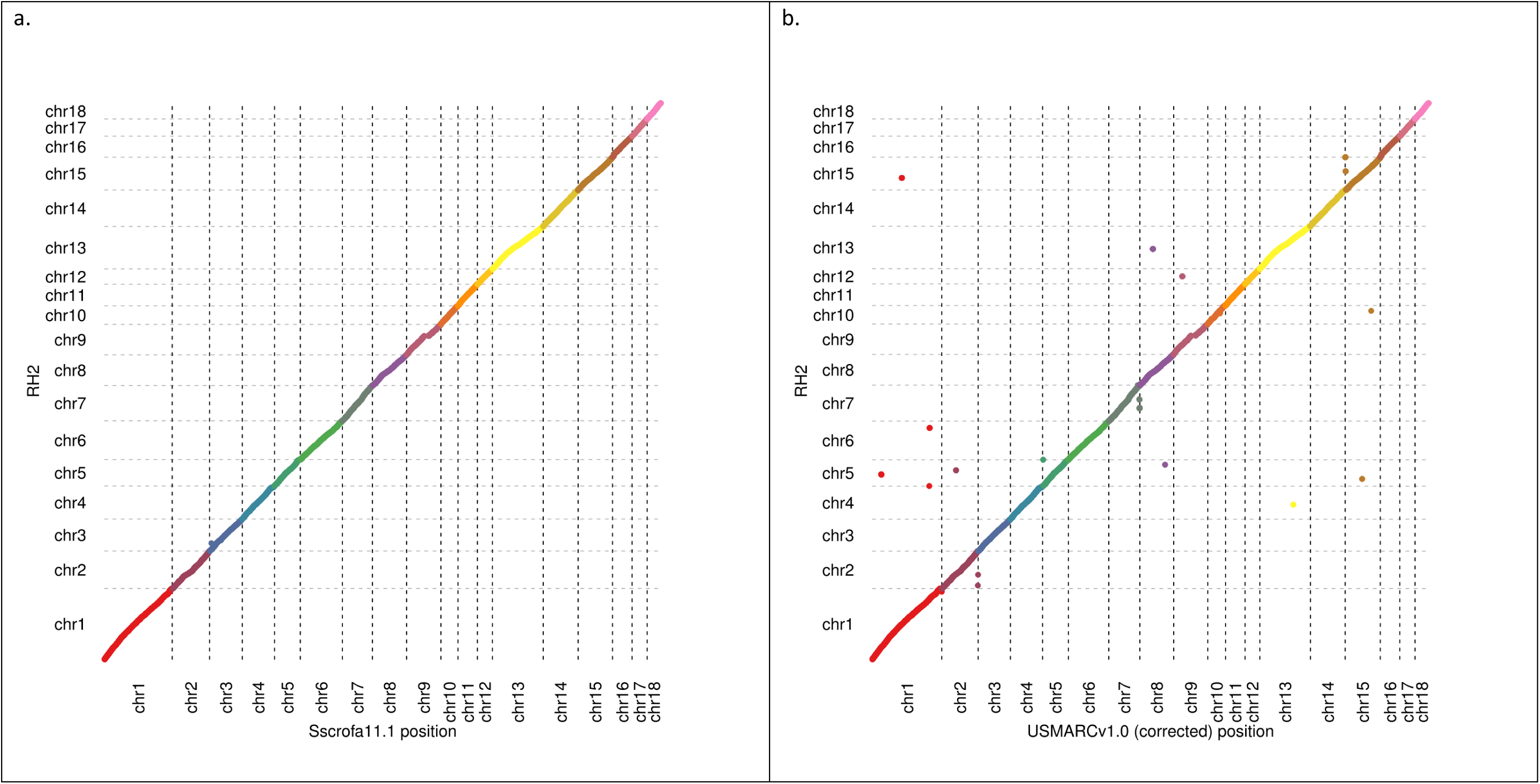
Assemblies and radiation hybrid map alignments. Plots illustrating co-linearity between radiation hybrid map and a) Sscrofa11.1 and b) USMARCv1.0 assemblies (autosomes only).

The quality of the Sscrofa11 assembly, which corresponds to Sscrofa11.1 after the exclusion of SSCY, was assessed as described previously for the existing Sanger sequence based draft assembly (Sscrofa10.2) [*14*]. Alignments of Illumina sequence reads from the same female pig were used to identify regions of low quality (LQ) or low coverage (LC) (Table 2). The analysis confirms that Sscrofa11 represents a significant improvement over the Sscrofa10.2 draft assembly. For example, the Low Quality Low Coverage (LQLC) proportion of the genome sequence has dropped from 33.07% to 16.3% when repetitive sequence is not masked, and falls to 1.6% when repeats are masked prior to read alignment. The remaining LQLC segments of Sscrofa11 may represent regions where short read coverage is low due to known systematic errors of the short read platform related to GC content, rather than deficiencies of the assembly.

**Table 2.** Summary of quality statistics for SSC1-18, SSCX. Quality measures and terms as defined [*14*].

|  | Mean<br>(Sscrofa11) | Std<br>(Sscrofa11) | Bases<br>(Sscrofa11) | % genome<br>(Sscrofa11) | % genome<br>(Sscrofa10.2) |
| --- | --- | --- | --- | --- | --- |
| High Coverage | 50 | 7 | 119,341,205 | 4.9 | 2.6 |
| Low Coverage (LC) | 50 | 7 | 185,385,536 | 7.5 | 26.6 |
| % Properly paired | 86 | 6.8 | 95,508,007 | 3.9 | 4.95 |
| % High inserts | 0.3 | 1.6 | 40,835,320 | 1.72 | 1.52 |
| % Low inserts | 8.2 | 4.3 | 114,793,298 | 4.7 | 3.99 |
| Low quality (LQ) | - | - | 284,838,040 | 11.6 | 13.85 |
| Total LQLC | - | - | 399,927,747 | 16.3 | 33.07 |
| LQLC windows that do not intersect RepeatMasker regions |  |  | 39,918,551 | 1.6 |  |

The Sscrofa11.1 assembly was also assessed visually using gEVAL [*29*]. The improvement in short range order and orientation as revealed by alignments with isogenic BAC and fosmid end sequences is illustrated for a particularly poor region of Sscrofa10.2 on chromosome 12 (Fig. S4). The problems in this area of Sscrofa10.2 arose from failures to order and orient the sequence contigs and resolve the redundancies between these sequence contigs within BAC clone CH242-147O24 (FP102566.2). The improved contiguity in Sscrofa11.1 not only resolves these local order and orientation errors, but also facilitates the annotation of a complete gene model for the *ABR* locus. Further examples of comparisons of Sscrofa10.2 and Sscrofa11.1 reveal improvements in contiguity, local order and orientation and gene models (Fig. S5 to S7).

### USMARCv1.0 assembly

Approximately sixty-five fold coverage of the genome of the MARC1423004 barrow was generated on a PacBio RSII instrument. The sequence was collected during the transition from P5/C3 to P6/C4 chemistry, with approximately equal numbers of subreads from each chemistry. A total of 199 cells of P5/C3 chemistry produced 95.3 Gb of sequence with mean subread length of 5.1 kb and subread N50 of 8.2 kb. A total of 127 cells of P6/C4 chemistry produced 91.6 Gb of sequence with mean subread length 6.5 kb and subread N50 of 10.3 kb, resulting in an overall average subread length, including data from both chemistries, of 6.4 kb. The reads were assembled using Celera Assembler 8.3rc2 [*30*] and Falcon (https://pb-falcon.readthedocs.io/en/latest/about.html). The resulting assemblies were compared and the Celera Assembler result was selected based on better agreement with a Dovetail Chicago® library [*25*], and was used to create a scaffolded assembly with the HiRise™ scaffolder consisting of 14,818 contigs with a contig N50 of 6.372 Mb (GenBank accession GCA_002844635.1; Table 1). The USMARCv1.0 scaffolds were therefore completely independent of the existing Sscrofa10.2 or new Sscrofa11.1 assemblies, and they can act as supporting evidence where they agree with those assemblies. However, chromosome assignment of the scaffolds was performed by alignment to Sscrofa10.2, and does not constitute independent confirmation of this ordering. The assignment of these scaffolds to individual chromosomes was confirmed post-hoc by FISH analysis as described for Sscrofa11.1 above. The FISH analysis revealed that several of these chromosome assemblies (SSC1, 5, 6-11, 13-16) are inverted with respect to the cytogenetic convention for pig chromosome (Table S4; Figs. S3, S8 to S10). After correcting the orientation of these inverted scaffolds, there is good agreement between the USMARCv1.0 assembly and the RH map [*9*] (Fig. 1b).

### Sscrofa11.1 and USMARCv1.0 are co-linear

The alignment of the two PacBio assemblies reveals a high degree of agreement and co-linearity, after correcting the inversions of several USMARCv1.0 chromosome assemblies (Fig. S11). The agreement between the Sscrofa11.1 and USMARCv1.0 assemblies is also evident in comparisons of specific loci (Figs. S5 to S7) although with some differences (e.g. Fig. S6). The whole genome alignment of Sscrofa11.1 and USMARCv1.0 (Fig. S11) masks some inconsistencies that are evident when the alignments are viewed on a single chromosome-by-chromosome basis (Figs. S8 to S10). It remains to be determined whether the small differences between the assemblies represent errors in the assemblies, or true structural variation between the two individuals (see discussion of the *ERLIN1* locus below).

Pairwise comparisons amongst the Sscrofa10.2, Sscrofa11.1 and USMARCv1.0 assemblies using the Assemblytics tools [*31*] (http://assemblytics.com) revealed a peak of insertions and deletion with sizes of about 300 bp (Figs. S12a to S12c). We assume that these correspond to SINE elements. Despite the fact that the Sscrofa10.2 and Sscrofa11.1 assemblies are representations of the same pig genome, there are many more differences between these assemblies than between the Sscrofa11.1 and USMARCv1.0 assemblies. We conclude that many of the differences between the Sscrofa11.1 assembly and the earlier Sscrofa10.2 assemblies represent improvements in the former. Some of the differences may indicate local differences in terms of which of the two haploid genomes has been captured in the assembly. The differences between the Sscrofa11.1 and USMARCv1.0 will represent a mix of true structural differences and assembly errors that will require further research to resolve. The Sscrofa11.1 and USMARCv1.0 assemblies were also compared to 11 Illumina short read assemblies [*17*] (Table S5a, b, c).

### Repetitive sequences, centromeres and telomeres

The repetitive sequence content of the Sscrofa11.1 and USMARCv1.0 was identified and characterised. These analyses allowed the identification of centromeres and telomeres for several chromosomes. The previous reference genome (Sscrofa10.2) that was established from Sanger sequence data and a minipig genome (minipig_v1.0, GCA_000325925.2) that was established from Illumina short read sequence data were also included for comparison. The numbers of the different repeat classes and the average mapped lengths of the repetitive elements identified in these four pig genome assemblies are summarised in Figures S13 and S14, respectively.

Putative telomeres were identified at the proximal ends of Sscrofa11.1 chromosome assemblies of SSC2, SSC3, SSC6, SSC8, SSC9, SSC14, SSC15, SSC18 and SSCX (Fig S1; Table S2). Putative centromeres were identified in the expected locations in the Sscrofa11.1 chromosome assemblies for SSC1-7, SSC9, SSC13 and SSC18 (Fig S2, Table S3). For the chromosome assemblies of each of SSC8, SSC11 and SSC15 two regions harbouring centromeric repeats were identified. Pig chromosomes SSC1-12 plus SSCX and SSCY are all metacentric, whilst chromosomes SSC13-18 are acrocentric. The putative centromeric repeats on SSC17 do not map to the expected end of the chromosome assembly.

### Completeness of the assemblies

The Sscrofa11.1 and USMARCv1.0 assemblies were assessed for completeness using two tools, BUSCO (Benchmarking Universal Single-Copy Orthologs) [*32*] and Cogent (https://github.com/Magdoll/Cogent). BUSCO uses a database of expected gene content based on near-universal single-copy orthologs from species with genomic data, while Cogent uses transcriptome data from the organism being sequenced, and therefore provides an organism-specific view of genome completeness. BUSCO analysis suggests both new assemblies are highly complete, with 93.8% and 93.1% of BUSCOs complete for Sscrofa11.1 and USMARCv1.0 respectively, a marked improvement on the 80.9% complete in Sscrofa10.2 and comparable to the human and mouse reference genome assemblies (Table S6).

Cogent is a tool that identifies gene families and reconstructs the coding genome using full-length, high-quality (HQ) transcriptome data without a reference genome and can be used to check assemblies for the presence of these known coding sequences. PacBio transcriptome (Iso-Seq) data consisting of high-quality isoform sequences from 7 tissues (diaphragm, hypothalamus, liver, skeletal muscle (*longissimus dorsi*), small intestine, spleen and thymus) [*33*] from the pig whose DNA was used as the source for the USMARCv1.0 assembly were pooled together for Cogent analysis. Cogent partitioned 276,196 HQ isoform sequences into 30,628 gene families, of which 61% had at least 2 distinct transcript isoforms. Cogent then performed reconstruction on the 18,708 partitions. For each partition, Cogent attempts to reconstruct coding ‘contigs’ that represent the ordered concatenation of transcribed exons as supported by the isoform sequences. The reconstructed contigs were then mapped back to Sscrofa11.1 and contigs that could not be mapped or map to more than one position are individually examined. There were five genes that were present in the Iso-Seq data, but missing in the Sscrofa11.1 assembly. In each of these five cases, a Cogent partition (which consists of 2 or more transcript isoforms of the same gene, often from multiple tissues) exists in which the predicted transcript does not align back to Sscrofa11.1. NCBI-BLASTN of the isoforms from the partitions revealed them to have near perfect hits with existing annotations for *CHAMP1*, *ERLIN1*, *IL1RN*, *MB*, and *PSD4*.

*ERLIN1* is missing from its predicted location on SSC14 between *CHUK* and *CPN1* gene in Sscrofa11.1. There is good support for the Sscrofa11.1 assembly in the region from the BAC end sequence alignments suggesting this area may represent a true haplotype. Indeed, a copy number variant (CNV) nsv1302227 has been mapped to this location on SSC14 [*34*] and the *ERLIN1* gene sequences present in BAC clone CH242-513L2 (ENA: CT868715.3) were incorporated into the earlier Sscrofa10.2 assembly. However, an alternative haplotype containing *ERLIN1* was not found in any of the assembled contigs from Falcon and this will require further investigation. The *ERLIN1* locus is present on SSC14 in the USMARCv1.0 assembly (30,107,816-30,143,074; note the USMARCv1.0 assembly of SSC14 is inverted relative to Sscrofa11.1). Of eleven short read pig genome assemblies [*17*] that have been annotated with the Ensembl pipeline (Ensembl release 98, September 2019) *ERLIN1* sequences are present in the expected genomic context in all eleven genome assemblies. As the *ERLIN1* gene is located at the end of a contig in eight of these short read assemblies, it suggests that this region of the pig genome presents difficulties for sequencing and assembly and the absence of *ERLIN1* in the Sscrofa11.1 is more likely to be an assembly error.

The other 4 genes are annotated in neither Sscrofa10.2 nor Sscrofa11.1. Two of these genes, *IL1RN* and *PSD4*, are present in the original Falcon contigs, however they were trimmed off during the contig QC stage because of apparent abnormal Illumina, BAC and fosmid mapping in the region which was likely caused by the repetitive nature of their expected location on chromosome 3 where a gap is present. The *IL1RN* and *PSD4* genes are present in the USMARCv1.0, albeit their location is anomalous, and are also present in the 11 short read assemblies [*17*]. *CHAMP1* (ENSSSCG00070014091) is present in the USMARCv1.0 assembly in the sub-telomeric region of the q-arm, after correcting the inversion of the USMARCv1.0 scaffold and is also present in all 11 short read assemblies [*17*]. After correcting the orientation of the USMARCv1.0 chromosome 11 scaffold there is a small inversion of the distal 1.07 Mbp relative to the Sscrofa11.1 assembly; this region harbours the *CHAMP1* gene. The orientation of the Sscrofa11.1 chromosome 11 assembly in this region is consistent with the predictions of the human-pig comparative map [*35*]. The myoglobin gene (*MB*) is present in the expected location in the USMARCv1.0 assembly flanked by *RASD2* and *RBFOX2*. Partial *MB* sequences are present distal to *RBFOX2* on chromosome 5 in the Sscrofa11.1 assembly. As there is no gap here in the Sscrofa11.1 assembly it is likely that the incomplete *MB* is a result of a misassembly in this region. This interpretation is supported by a break in the pairs of BAC and fosmid end sequences that map to this region of the Sscrofa11.1 assembly. Some of the expected gene content missing from this region of the Sscrofa11.1 chromosome 5 assembly, including *RASD2*, *HMOX1* and *LARGE1* is present on an unplaced scaffold (AEMK02000361.1). Cogent analysis also identified 2 cases of potential fragmentation in the Sscrofa11.1 genome assembly that resulted in the isoforms being mapped to two separate loci, though these will require further investigation. In summary, the BUSCO and Cogent analyses indicate that the Sscrofa11.1 assembly captures a very high proportion of the expressed elements of the genome.

### Improved annotation

Annotation of Sscrofa11.1 was carried out with the Ensembl annotation pipeline and released via the Ensembl Genome Browser [*36*] (http://www.ensembl.org/Sus_scrofa/Info/Index) (Ensembl release 90, August 2017). Statistics for the annotation as updated in June 2019 (Ensembl release 98, September 2019) are listed in Table 3. This annotation is more complete than that of Sscrofa10.2 and includes fewer fragmented genes and pseudogenes.

**Table 3.** Annotation statistics. Ensembl annotation of pig (Sscrofa10.2, Sscrofa11.1, USMARCv1.0), human (GRCh38.p12) and mouse (GRCm38.p6) assemblies.

|  | Sscrofa10.2 | Sscrofa11.1 | USMARCv1.0 | GRCh38.p13 | GRCm38.p6 |
| --- | --- | --- | --- | --- | --- |
|  | Ensembl (Release 89) | Ensembl (Release 98) | Ensembl (Release 97) | Ensembl (Release 98) | Ensembl (Release 98) |
| Coding genes | 21,630<br>(Incl. 10 read<br>through) | 21,301 | 21,535 | 20,444<br>incl 667 read through | 22,508<br>incl 270 read through |
| Non-coding genes | 3,124 | 8,971 | 6,113 | 23,949 | 16,078 |
| small non-coding genes | 2,804 | 2,156 | 2,427 | 4,871 | 5,531 |
| long non-coding genes | 135<br>(incl 1 read through) | 6,798 | 3,307 | 16,857<br>incl 304 read through | 9,985<br>incl 75 read through |
| misc. non-coding genes | 185 | 17 | 379 | 2,221 | 562 |
| Pseudogenes | 568 | 1,626 | 674 | 15,214<br>incl 8 read through | 13,597<br>incl 4 read through |
| Gene transcripts | 30,585 | 63,041 | 58,692 | 227,530 | 142,446 |
| Genscan gene<br>predictions | 52,372 | 46,573 | 152,168 | 51,756 | 57,381 |
| Short variants | 60,389,665 | 64,310,125 |  | 665,834,144 | 83,761,978 |
| Structural variants | 224,038 | 224,038 |  | 6,013,113 | 791,878 |

The annotation pipeline utilised extensive short read RNA-Seq data from 27 tissues and long read PacBio Iso-Seq data from 9 adult tissues. This provided an unprecedented window into the pig transcriptome and allowed for not only an improvement to the main gene set, but also the generation of tissue-specific gene tracks from each tissue sample. The use of Iso-Seq data also improved the annotation of UTRs, as they represent transcripts sequenced across their full length from the polyA tract.

In addition to improved gene models, annotation of the Sscrofa11.1 assembly provides a more complete view of the porcine transcriptome than annotation of the previous assembly (Sscrofa10.2; Ensembl releases 67-89, May 2012 – May 2017) with increases in the numbers of transcripts annotated (Table 3). However, the number of annotated transcripts remains lower than in the human and mouse genomes. The annotation of the human and mouse genomes and in particular the gene content and encoded transcripts has been more thorough as a result of extensive manual annotation.

Efforts were made to annotate important classes of genes, in particular immunoglobulins and olfactory receptors. For these genes, sequences were downloaded from specialist databases and the literature in order to capture as much detail as possible (see supplementary information for more details).

These improvements in terms of the resulting annotation were evident in the results of the comparative genomics analyses run on the gene set. The previous annotation had 12,919 one-to-one orthologs with human, while the new annotation of the Sscrofa11.1 assembly has 15,544. Similarly, in terms of conservation of synteny, the previous annotation had 11,661 genes with high confidence gene order conservation scores, while the new annotation has 15,958. There was also a large reduction in terms of genes that were either abnormally short or split when compared to their orthologs in the new annotation.

The Sscrofa11.1 assembly has also been annotated using the NCBI pipeline (https://www.ncbi.nlm.nih.gov/genome/annotation_euk/Sus_scrofa/106/). We have compared these two annotations. The Ensembl and NCBI annotations of Sscrofa11.1 are broadly similar (Table S7). There are 17,676 protein coding genes and 1,700 non-coding genes in common. However, 540 of the genes annotated as protein-coding by Ensembl are annotated as non-coding or pseudogenes by NCBI and 227 genes annotated as non-coding by NCBI are annotated as protein-coding (215) or as pseudogenes (12) by Ensembl. The NCBI RefSeq annotation can be visualised in the Ensembl Genome Browser by loading the RefSeq GFF3 track and the annotations compared at the individual locus level. Similarly, the Ensembl annotated genes can be visualised in the NCBI Genome Browser. Despite considerable investment there are also differences in the Ensembl and NCBI annotation of the human reference genome sequence with 20,444 and 19,755 protein-coding genes on the primary assembly, respectively. The MANE (Matched Annotation from NCBI and EMBL-EBI) project was launched to resolve these differences and identify a matched representative transcript for each human protein-coding gene (https://www.ensembl.org/info/genome/genebuild/mane.html). To date a MANE transcript has been identified for 12,985 genes.

We have also annotated the USMARCv1.0 assembly using the Ensembl pipeline [*36*] and this annotation was released via the Ensembl Genome Browser (https://www.ensembl.org/Sus_scrofa_usmarc/Info/Index) (Ensembl release 97, July 2019; see Table 3 for summary statistics). More recently, we have annotated a further eleven short read pig genome assemblies [*17*] (Ensembl release 98, September 2019, see Tables S5c and S10 for summary statistics for the assemblies and annotation, respectively).

### SNP chip probes mapped to assemblies

The probes from four commercial SNP chips were mapped to the Sscrofa10.2, Sscrofa11.1 and USMARCv1.0 assemblies. We identified 1,709, 56, and 224 markers on the PorcineSNP60, GGP LD and 80K commercial chips that were previously unmapped and now have coordinates on the Sscrofa11.1 reference (Table S8). These newly mapped markers can now be imputed into a cross-platform, common set of SNP markers for use in genomic selection. Additionally, we have identified areas of the genome that are poorly tracked by the current set of commercial SNP markers. The previous Sscrofa10.2 reference had an average marker spacing of 3.57 kbp (Stdev: 26.5 kb) with markers from four commercial genotyping arrays. We found this to be an underestimate of the actual distance between markers, as the Sscrofa11.1 reference coordinates consisted of an average of 3.91 kbp (Stdev: 14.9 kbp) between the same set of markers. We also found a region of 2.56 Mbp that is currently devoid of suitable markers on the new reference.

A Spearman’s rank order (rho) value was calculated for each assembly (alternative hypothesis: rho is equal to zero; p < 2.2 x 10^−16^): Sscrofa10.2: 0.88464; Sscrofa11.1: 0.88890; USMARCv1.0: 0.81260. This rank order comparison was estimated by ordering all of the SNP probes from all chips by their listed manifest coordinates against their relative order in each assembly (with chromosomes ordered by karyotype). Any unmapped markers in an assembly were penalized by giving the marker a “−1” rank in the assembly ranking order.

In order to examine general linear order of placed markers on each assembly, the marker rank order (y axis; used above in the Spearman’s rank order test) was plotted against the rank order of the probe rank order on the manifest file (x axis) (Fig. S15). The analyses revealed some interesting artefacts that suggest that the SNP manifest coordinates for the porcine 60K SNP chip are still derived from an obsolete (Sscrofa9) reference in contrast to all other manifests (Sscrofa10.2). Also, it confirms that several of the USMARCv1.0 chromosome scaffolds are inverted with respect to the canonical orientation of pig chromosomes. The large band of points at the top of the plot corresponds to marker mappings on the unplaced contigs of each assembly. These unplaced contigs often correspond to assemblies of alternative haplotypes in heterozygous regions of the reference animal [*24*]. Marker placement on these segments suggests that these variants are tracking different haplotypes in the population, which is the desired intent of genetic markers used in Genomic Selection.

## Discussion

We have assembled a superior, extremely continuous reference assembly (Sscrofa11.1) by leveraging the excellent contig lengths provided by long reads, and a wealth of available data including Illumina paired-end, BAC end sequence, finished BAC sequence, fosmid end sequences, and the earlier curated draft assembly (Sscrofa10.2). The pig genome assemblies USMARCv1.0 and Sscrofa11.1 reported here are 92-fold to 694-fold respectively, more continuous than the published draft reference genome sequence (Sscrofa10.2) [*13*]. The new pig reference genome assembly (Sscrofa11.1) with its contig N50 of 48,231,277 bp and 506 gaps compares favourably with the current human reference genome sequence (GRCh38.p12) that has a contig N50 of 57,879,411 bp and 875 gaps (Table 1). Indeed, considering only the chromosome assemblies built on PacBio long read data (i.e. Sscrofa11 - the autosomes SSC1-SSC18 plus SSCX), there are fewer gaps in the pig assembly than in human reference autosomes and HSAX assemblies. Most of the gaps in the Sscrofa11.1 reference assembly are attributed to the fragmented assembly of SSCY. The capturing of centromeres and telomeres for several chromosomes (Tables S2, S3; Figs. S1, S2) provides further evidence that the Sscrofa11.1 assembly is more complete. The increased contiguity of Sscrofa11.1 is evident in the graphical comparison to Sscrofa10.2 illustrated in Figure 2.

**Figure 2.**
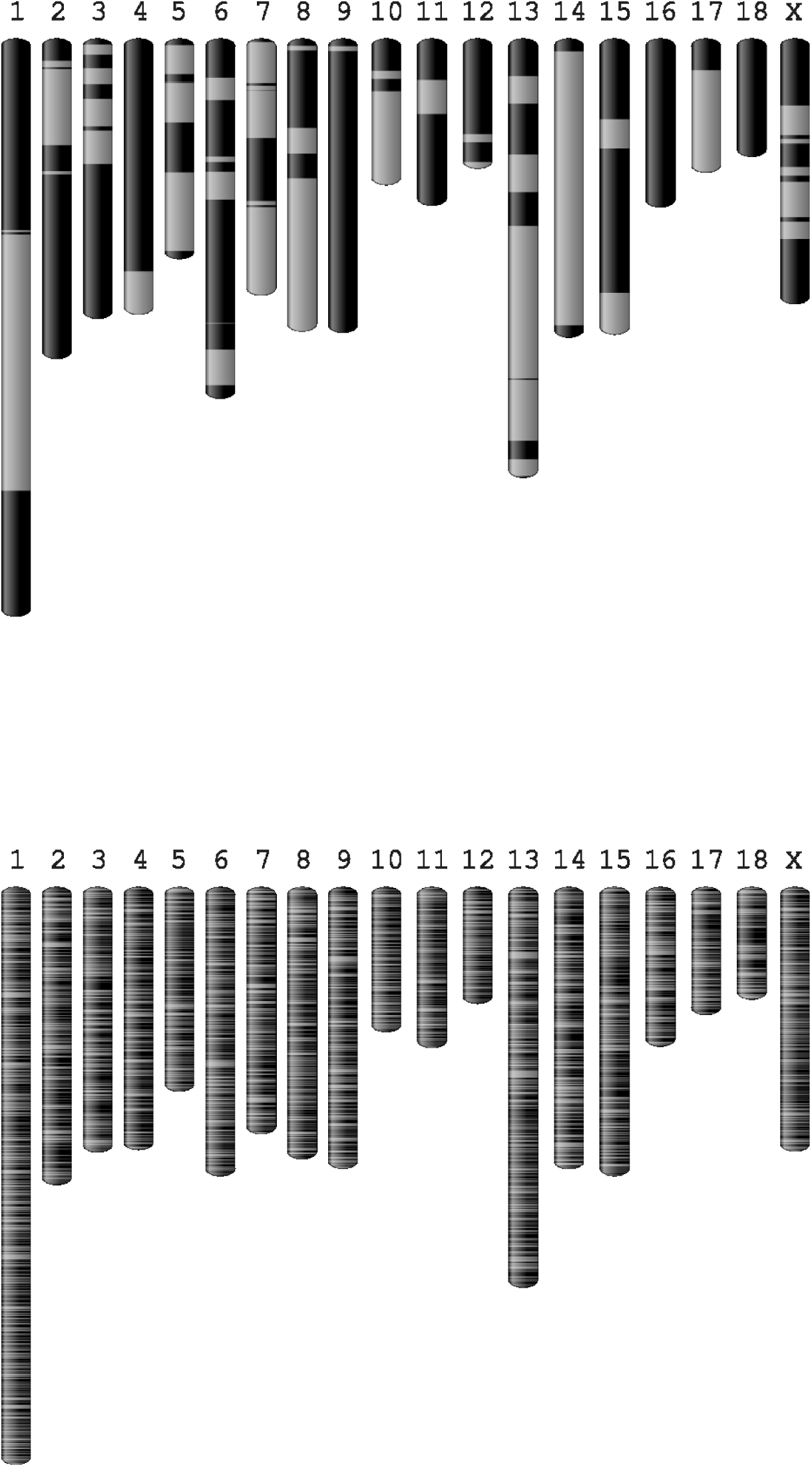
Visualisation of improvements in assembly contiguity. Graphical visualisation of contigs for Sscrofa11 (top) and Sscrofa10.2 (bottom) as alternating dark and light grey bars.

The improvements in the reference genome sequence (Sscrofa11.1) relative to the draft assembly (Sscrofa10.2) [*13*] are not restricted to greater continuity and fewer gaps. The major flaws in the BAC clone-based draft assembly were i) failures to resolve the sequence redundancy amongst sequence contigs within BAC clones and between adjacent overlapping BAC clones and ii) failures to accurately order and orient the sequence contigs within BAC clones. Although the Sanger sequencing technology used has a much lower raw error rate than the PacBio technology, the sequence coverage was only 4-6 fold across the genome. The improvements in continuity and quality (Table 2; Figs. S5 to S7) have yielded a better template for annotation resulting in better gene models. The Sscrofa11.1 and USMARCv1.0 assemblies are classed as 4|4|1 and 3|5|1 [10^X^: N50 contig (kb); 10^Y^: N50 scaffold (kb); Z = 1|0: assembled to chromosome level] respectively compared to Sscrofa10.2 as 1|2|1 and the human GRCh38p5 assembly as 4|4|1 (see https://geval.sanger.ac.uk).

The improvement in the complete BUSCO (Benchmarking Universal Single-Copy Orthologs) genes indicates that both Sscrofa11.1 and USMARCv1.0 represent superior templates for annotation of gene models than the draft Sscrofa10.2 assembly and are comparable to the finished human and mouse reference genome sequences (Table S6). Further, a companion bioinformatics analysis of available Iso-seq and companion Ilumina RNA-seq data across the nine tissues surveyed has identified a large number (>54,000) of novel transcripts [*33*]. A majority of these transcripts are predicted to be spliced and validated by RNA-seq data. Beiki and colleagues identified 10,465 genes expressing Iso-seq transcripts that are present on the Sscrofa11.1 assembly, but which are unannotated in current NCBI or Ensembl annotations.

Whilst the alignment of the Sscrofa11.1 and USMARCv1.0 assemblies revealed that several of the USMARCv1.0 chromosome assemblies are inverted relative to Sscrofa11.1 and the cytogenetic map. Such inversions are due to the agnostic nature of genome assembly and post-assembly polishing programs. Unless these are corrected post-hoc by manual curation, they result in artefactual inversions of the entire chromosome. However, such inversions do not generally impact downstream analysis that does not involve the relative order/orientation of whole chromosomes.

Whether the differences between Sscrofa11.1 and USMARCv1.0 in order and orientation within chromosomes represent assembly errors or real chromosomal differences will require further research. The sequence present at the telomeric end of the long arm of the USMARCv1.0 chromosome 7 assembly (after correcting the orientation of the USMARCv1.0 SSC7) is missing from the Sscrofa11.1 SSC7 assembly, and currently located on a 3.8 Mbp unplaced scaffold (AEMK02000452.1). This unplaced scaffold harbours several genes including *DIO3*, *CKB* and *NUDT14* whose orthologues map to human chromosome 14 as would be predicted from the pig-human comparative map [*35*]. This omission will be corrected in an updated assembly in future.

We demonstrate moderate improvements in the placement and ordering of commercial SNP genotyping markers on the Sscrofa11.1 reference genome which will impact future genomic selection programs. The reference-derived order of SNP markers plays a significant role in imputation accuracy, as demonstrated by a whole-genome survey of misassembled regions in cattle that found a correlation between imputation errors and misassemblies [*37*]. The gaps in SNP chip marker coverage that we identified will inform future marker selection surveys, which are likely to prioritize regions of the genome that are not currently being tracked by marker variants in close proximity to potential causal variant sites. In addition to the gaps in coverage provided by the commercial SNP chips there are regions of the genome assemblies that are devoid of annotated sequence variation as hitherto sequence variants have been discovered against incomplete genome assemblies. Thus, there is a need to re-analyse good quality re-sequence data against the new assemblies in order to provide a better picture of sequence variation in the pig genome.

The cost of high coverage whole-genome sequencing (WGS) precludes it from routine use in breeding programs. However, it has been suggested that low coverage WGS followed by imputation of haplotypes may be a cost-effective replacement for SNP arrays in genomic selection [*38*]. Imputation from low coverage sequence data to whole genome information has been shown to be highly accurate [*39, 40*]. At the 2018 World Congress on Genetics Applied to Livestock Production Aniek Bouwman reported that in a comparison of Sscrofa10.2 with Sscrofa11.1 (for SSC7 only) for imputation from 600K SNP genotypes to whole genome sequence overall imputation accuracy on SSC7 improved considerably from 0.81 (1,019,754 variants) to 0.90 (1,129,045 variants) (Aniek Bouwman, pers. comm). Thus, the improved assembly may not only serve as a better template for discovering genetic variation but also have advantages for genomic selection, including improved imputation accuracy.

Advances in the performance of long read sequencing and scaffolding technologies, improvements in methods for assembling the sequence reads and reductions in costs are enabling the acquisition of ever more complete genome sequences for multiple species and multiple individuals within a species. For example, in terms of adding species, the Vertebrate Genomes Project (https://vertebrategenomesproject.org/) aims to generate error-free, near gapless, chromosomal level, haplotyped phase assemblies of all of the approximately 66,000 vertebrate species and is currently in its first phase that will see such assemblies created for an exemplar species from all 260 vertebrate orders. At the level of individuals within a species, smarter assembly algorithms and sequencing strategies are enabling the production of high quality truly haploid genome sequences for outbred individuals [*24*]. The establishment of assembled genome sequences for key individuals in the nucleus populations of the leading pig breeding companies is achievable and potentially affordable. However, 10-30x genome coverage short read data generated on the Illumina platform and aligned to a single reference genome is likely to remain the primary approach to sequencing multiple individuals within farmed animal species such as cattle and pigs [*21, 41*].

There are significant challenges in making multiple assembled genome resources useful and accessible. The current paradigm of presenting a reference genome as a linear representation of a haploid genome of a single individual is an inadequate reference for a species. As an interim solution the Ensembl team are annotating multiple assemblies for some species such as mouse (https://www.ensembl.org/Mus_musculus/Info/Strains) [*42*]. We have implemented this solution for pig genomes, including eleven Illumina short-read assemblies [*17*] in addition to the reference Sscrofa11.1 and USMARCv1.0 assemblies reported here (Ensembl release 98, September 2019 https://www.ensembl.org/Sus_scrofa/Info/Strains?db=core). Although these additional pig genomes are highly fragmented (Table S5c) with contig N50 values from 32 – 102 kbp, the genome annotation (Table S10) provides a resource to explore pig gene space across thirteen genomes, including six Asian pig genomes. The latter are important given the deep phylogenetic split of about 1 million years between European and Asian pigs [*13*].

The current human genome reference already contains several hundred alternative haplotypes and it is expected that the single linear reference genome of a species will be replaced with a new model – the graph genome [*43–45*]. These paradigm shifts in the representation of genomes present challenges for current sequence alignment tools and the ‘best-in-genome’ annotations generated thus far. The generation of high quality annotation remains a labour-intensive and time-consuming enterprise. Comparisons with the human and mouse reference genome sequences which have benefited from extensive manual annotation indicate that there is further complexity in the porcine genome as yet unannotated (Table 3). It is very likely that there are many more transcripts, pseudogenes and non-coding genes (especially long non-coding genes), to be discovered and annotated on the pig genome sequence [*33*]. The more highly continuous pig genome sequences reported here provide an improved framework against which to discover functional sequences, both coding and regulatory, and sequence variation. After correction for some contig/scaffold inversions in the USMARCv1.0 assembly, the overall agreement between the assemblies is high and illustrates that the majority of genomic variation is at smaller scales of structural variation. However, both assemblies still represent a composite of the two parental genomes present in the animals, with unknown effects of haplotype switching on the local accuracy across the assembly.

Future developments in high quality genome sequences for the domestic pig are likely to include: (i) gap closure of Sscrofa11.1 to yield an assembly with one contig per (autosomal) chromosome arm exploiting the isogenic BAC and fosmid clone resource as illustrated here for chromosome 16 and 18; and (ii) haplotype resolved assemblies of a Meishan and White Composite F1 crossbred pig currently being sequenced. Beyond this haplotype resolved assemblies for key genotypes in the leading pig breeding company nucleus populations and of miniature pig lines used in biomedical research can be anticipated in the next 5 years. Unfortunately, some of these genomes may not be released into the public domain. The first wave of results from the Functional Annotation of ANimal Genomes (FAANG) initiative [*46, 47*], are emerging and will add to the richness of pig genome annotation.

In conclusion, the new pig reference genome (Sscrofa11.1) described here represents a significantly enhanced resource for genetics and genomics research and applications for a species of importance to agriculture and biomedical research.

## Methods

Additional detailed methods and information on the assemblies and annotation are included in the Supplementary Materials.

### Preparation of genomic DNA

DNA was extracted from Duroc 2-14 cultured fibroblast cells passage 16-18 using the Qiagen Blood & Cell Culture DNA Maxi Kit. DNA was isolated from lung tissue from barrow MARC1423004 using a salt extraction method.

### Genome sequencing and assembly

Genomic DNAs from the samples described above were used to prepare libraries for sequencing on Pacific Biosciences RS II sequencer [*48*]. For Duroc 2-14 DNA P6/C4 chemistry was used, whilst for MARC1423004 DNA a mix of P6/C4 and earlier P5/C3 chemistry was used.

Reads from the Duroc 2-14 DNA were assembled into contigs using the Falcon v0.4.0 assembly pipeline following the standard protocol [*26*]. Quiver v. 2.3.0 [*49*] was used to correct the primary and alternative contigs. Only the primary pseudo-haplotype contigs were used in the assembly. The reads from the MARC1423004 DNA were assembled into contigs using Celera Assembler v8.3rc2 [*30*]. The contigs were scaffolded as described in the results section above.

### Fluorescence *in situ* hybridisation

Metaphase preparations were fixed to slides and dehydrated through an ethanol series (2 min each in 2×SSC, 70%, 85% and 100% ethanol at RT). Probes were diluted in a formamide buffer (Cytocell) with Porcine Hybloc (Insight Biotech) and applied to the metaphase preparations on a 37°C hotplate before sealing with rubber cement. Probe and target DNA were simultaneously denatured for 2 mins on a 75°C hotplate prior to hybridisation in a humidified chamber at 37°C for 16 h. Slides were washed post hybridisation in 0.4x SSC at 72°C for 2 mins followed by 2x SSC/0.05% Tween 20 at RT for 30 secs, and then counterstained using VECTASHIELD anti-fade medium with DAPI (Vector Labs). Images were captured using an Olympus BX61 epifluorescence microscope with cooled CCD camera and SmartCapture (Digital Scientific UK) system.

### Analysis of repetitive sequences, including telomeres and centromeres

Repeats were identified using RepeatMasker (v.4.0.7) (http://www.repeatmasker.org) with a combined repeat database including Dfam (v.20170127) [*50*] and RepBase (v.20170127) [*51*]. RepeatMasker was run with “sensitive” (-s) setting using sus scrofa as the query species (-- species “sus scrofa”). Repeats which showed greater than 40% sequence divergence or were shorter than 70% of the expected sequence length were filtered out from subsequent analyses. The presence of potentially novel repeats was assessed by RepeatMasker using the novel repeat library generated by RepeatModeler (v.1.0.11) (http://www.repeatmasker.org).

Telomeres were identified by running Tandem Repeat Finder (TRF) [*52*] with default parameters apart from Mismatch (5) and Minscore (40). The identified repeat sequences were then searched for the occurrence of five identical, consecutive units of the TTAGGG vertebrate motif or its reverse complement and total occurrences of this motif was counted within the tandem repeat. Regions which contained at least 200 identical hexamer units, were >2kb of length and had a hexamer density of >0.5 were retained as potential telomeres.

Centromeres were predicted using the following strategy. First, the RepeatMasker output, both default and novel, was searched for centromeric repeat occurrences. Second, the assemblies were searched for known, experimentally verified, centromere specific repeats [*53, 54*] in the Sscrofa11.1 genome. Then the three sets of repeat annotations were merged together with BEDTools [*55*] (median and mean length: 786 bp and 5775 bp, respectively) and putative centromeric regions closer than 500 bp were collapsed into longer super-regions. Regions which were >5kb were retained as potential centromeric sites.

### Long read RNA sequencing (Iso-Seq)

The following tissues were harvested from MARC1423004 at age 48 days: brain (BioSamples: SAMN05952594), diaphragm (SAMN05952614), hypothalamus (SAMN05952595), liver (SAMN05952612), small intestine (SAMN05952615), skeletal muscle – *longissimus dorsi* (SAMN05952593), spleen (SAMN05952596), pituitary (SAMN05952626) and thymus (SAMN05952613). Total RNA from each of these tissues was extracted using Trizol reagent (ThermoFisher Scientific) and the provided protocol. Briefly, approximately 100 mg of tissue was ground in a mortar and pestle cooled with liquid nitrogen, and the powder was transferred to a tube with 1 ml of Trizol reagent added and mixed by vortexing. After 5 minutes at room temperature, 0.2 mL of chloroform was added and the mixture was shaken for 15 seconds and left to stand another 3 minutes at room temperature. The tube was centrifuged at 12,000 x g for 15 minutes at 4°C. The RNA was precipitated from the aqueous phase with 0.5 mL of isopropanol. The RNA was further purified with extended DNase I digestion to remove potential DNA contamination. The RNA quality was assessed with a Fragment Analyzer (Advanced Analytical Technologies Inc., IA). Only RNA samples of RQN above 7.0 were used for library construction. PacBio IsoSeq libraries were constructed per the PacBio IsoSeq protocol. Briefly, starting with 3 μg of total RNA, cDNA was synthesized by using SMARTer PCR cDNA Synthesis Kit (Clontech, CA) according to the IsoSeq protocol (Pacific Biosciences, CA). Then the cDNA was amplified using KAPA HiFi DNA Polymerase (KAPA Biotechnologies) for 10 or 12 cycles followed by purification and size selection into 4 fractions: 0.8-2 kb, 2-3 kb, 3-5 kb and >5 kb. The fragment size distribution was validated on a Fragment Analyzer (Advanced Analytical Technologies Inc, IA) and quantified on a DS-11 FX fluorometer (DeNovix, DE). After a second round of large scale PCR amplification and end repair, SMRT bell adapters were separately ligated to the cDNA fragments. Each size fraction was sequenced on 4 or 5 SMRT Cells v3 using P6-C4 chemistry and 6 hour movies on a PacBio RS II sequencer (Pacific Bioscience, CA). Short read RNA-Seq libraries were also prepared for all nine tissue using TruSeq stranded mRNA LT kits and supplied protocol (Illumina, CA), and sequenced on a NextSeq500 platform using v2 sequencing chemistry to generate 2 x 75 bp paired-end reads.

The Read of Insert (ROI) were determined by using *consensustools.sh* in the SMRT-Analysis pipeline v2.0, with reads which were shorter than 300 bp and whose predicted accuracy was lower than 75% removed. Full-length, non-concatemer (FLNC) reads were identified by running the classify.py command. The cDNA primer sequences as well as the poly(A) tails were trimmed prior to further analysis. Paired-end Illumina RNA-Seq reads from each tissue sample were trimmed to remove the adaptor sequences and low-quality bases using Trimmomatic (v0.32) [*56*] with explicit option settings: *ILLUMINACLIP:adapters.fa: 2:30:10:1:true LEADING:3 TRAILING:3 SLIDINGWINDOW: 4:20 LEADING:3 TRAILING:3 MINLEN:25*, and overlapping paired-end reads were merged using the PEAR software (v0.9.6) [*57*]. Subsequently, the merged and unmerged RNA-Seq reads from the same tissue samples were *in silico* normalized in a mode for single-end reads by using a Trinity (v2.1.1) [*58*] utility*, insilico_read_normalization.pl*, with the following settings*: --max_cov 50 --max_pct_stdev 100 -- single*. Errors in the full-length, non-concatemer reads were corrected with the preprocessed RNA-Seq reads from the same tissue samples by using proovread (v2.12) [*59*]. Untrimmed sequences with at least some regions of high accuracy in the *.trimmed.fq* files were extracted based on sequence IDs in *.untrimmed.fa* files to balance off the contiguity and accuracy of the final reads.

### Short read RNA sequencing (RNA-Seq)

In addition to the Illumina short read RNA-seq data generated from MARC1423004 and used to correct the Iso-Seq data (see above), Illumina short read RNA-seq data (PRJEB19386) were also generated from a range of tissues from four juvenile Duroc pigs (two male, two female) and used for annotation as described below. Extensive metadata with links to the protocols for sample collection and processing are linked to the BioSample entries under the Study Accession PRJEB19386. The tissues sampled are listed in Table S9. Sequencing libraries were prepared using a ribodepletion TruSeq stranded RNA protocol and 150 bp paired end sequences generated on the Illumina HiSeq 2500 platform in rapid mode.

### Annotation

The assembled genomes were annotated using the Ensembl pipelines [*36*] as detailed in the Supplementary materials. The Iso-Seq and RNA-Seq data described above were used to build gene models.

### Mapping SNP chip probes

The probes from four commercial SNP chips were mapped to the Sscrofa10.2, Sscrofa11.1 and USMARCv1.0 assemblies using BWA MEM [*60*] and a wrapper script (https://github.com/njdbickhart/perl_toolchain/blob/master/assembly_scripts/alignAndOrderSnpProbes.pl). Probe sequence was derived from the marker manifest files that are available on the provider websites: Illumina PorcineSNP60 https://emea.illumina.com/products/by-type/microarray-kits/porcine-snp60.html) [*1*]; Affymetrix Axiom™ Porcine Genotyping Array (https://www.thermofisher.com/order/catalog/product/550588); Gene Seek Genomic Profiler Porcine – HD beadChip (http://genomics.neogen.com/uk/ggp-porcine); Gene Seek Genomic Profiler Porcine v2–LD Chip (http://genomics.neogen.com/uk/ggp-porcine). In order to retain marker manifest coordinate information, each probe marker name was annotated with the chromosome and position of the marker’s variant site from the manifest file. All mapping coordinates were tabulated into a single file, and were sorted by the chromosome and position of the manifest marker site. In order to derive and compare relative marker rank order, a custom Perl script (https://github.com/njdbickhart/perl_toolchain/blob/master/assembly_scripts/pigGenomeSNPSortRankOrder.pl) was used to sort and number markers based on their mapping locations in each assembly.

## Supplementary materials

Supplementary materials for this article include:

Supplementary Methods and Information

Table S1. Pacific Biosciences read statistics.

Table S2. Predicted telomeres.

Table S3. Predicted centromeres.

Table S4. Assigning scaffolds to chromosomes.

Table S5. Assemblytics comparisons.

Table S6. BUSCO results.

Table S7. Annotation statistics. (Ensembl-NCBI comparison)

Table S8. Commercial SNP chip probes.

Table S9. Tissue samples.

Table S10. Ensembl annotation statistics for 13 pig genome assemblies

Figure S1. Predicted telomeres.

Figure S2. Predicted centromeres.

Figure S3. Fluorescent *in situ* hybridisation assignments.

Fig. S4. Improvement in local order and orientation and reduction in redundancy.

Fig. S5. Assembly comparisons in gEVAL (SSC15).

Fig. S6. Assembly comparisons in gEVAL (SSC5).

Fig. S7. Assembly comparisons in gEVAL (SSC18).

Fig. S8. Order and orientation of SSC18 assemblies.

Fig. S9. Order and orientation of SSC7 assemblies.

Fig. S10. Order and orientation of SSC8 assemblies.

Fig. S11. Assembly alignments.

Figure S12. Assemblytics results.

Figure S13. Counts of repetitive elements in four pig assemblies.

Figure S14. Average mapped length of repetitive elements in four pig genomes.

Figure S15. Assembly SNP rank concordance versus reported chromosomal location.

## Acknowledgements

We are grateful to Chris Tyler-Smith (Wellcome Trust Sanger Institute) for sharing the SSCY sequence data for Sscrofa11.1.

## Author contributions

A.L.A. and T.P.L.S. conceived, coordinated and managed the project; A.L.A., P.F., D.A.H., T.P.L.S. M.W. supervised staff and students performing the analyses; D.J.N., L.R., L.B.S., T.P.L.S. provided biological resources; R.H., K.S.K. and T.P.L.S. generated PacBio sequence data; H.A.F., T.P.L.S. and R.T. generated Illumina WGS and RNA-Seq data; N.A.A., C.A.S., B.M.S. provided SSCY assemblies; D.J.N, and T.P.L.S. generated Iso-Seq data; G.H., R.H., S.K., A.M.P., A.S.S, A.W. generated sequence assemblies; A.W. polished and quality checked Sscrofa11.1; W.C., G.H., K.H., S.K., B.D.R., A.S.S., S.G.S., E.T. performed quality checks on the sequence assemblies; R.E.O’C. and D.K.G. performed cytogenetics analyses; L.E. analysed repeat sequences; H.B., H.L., N.M., C.K.T. analysed Iso-Seq data; D.M.B. and G.A.R. analysed sequence variants; B.A., K.B., C.G.G., T.H., O.I., F.J.M. annotated the assembled genome sequences; A.W. and A.L.A drafted the manuscript; all authors read and approved the final manuscript.

## Data and materials availability

The genome assemblies are deposited at NCBI under accession numbers GCA_000003025 (Sscrofa11.1) and GCA_002844635.1 (USMARCv1.0). The associated BioSample accession numbers are SAMN02953785 and SAMN07325927, respectively. Iso-seq and RNA-Seq data used for analysis and annotation are available under accession numbers PRJNA351265 and PRJEB19386, respectively.

