## Supplementary information for "An improved pig reference genome sequence to enable pig genetics and genomics research"

### On-line Methods and Supplementary information

### 1. Sequence and assembly

#### 1.1 From Duroc 2-14 DNA to Sscrofa11.1 assembly

##### 1.1.1 Sample, sequencing and assembly

DNA was extracted from Duroc 2-14 cultured fibroblast cells passage 16-18 using the Qiagen Blood & Cell Culture DNA Maxi Kit, producing 139.15 µg DNA from three extractions. The high molecular weight DNA from this extraction was sequenced by Pacific Biosciences (PacBio) using their long read sequencing technology. Libraries for SMRT sequencing were prepared and sequenced as described previously [*48*] using P6-C4 chemistry on the RSII using 213 SMRT cells.

Contigs were assembled using the Falcon v0.4.0 assembly pipeline following the standard protocol. Quiver v. 2.3.0 [*49*] was used to correct the primary and alternative contigs. Only the primary pseudo-haplotype contigs were used in the assembly.

##### 1.1.2 Contig quality assessment and contig splitting

Paired-end Illumina reads from the same individual (http://www.ebi.ac.uk/ena/data/view/PRJEB9115) were mapped to the 3,206 haploid contigs and assessed for structural abnormalities using the methods described previously [*14*]. Briefly, 1,000 bp windows across the contigs were assessed for levels of abnormal mapping including high GC-normalized coverage, improper pairing and unexpected insert sizes. Additionally BAC end sequences (BES) (CHORI-242 library) [*12*] and fosmids (WTSI_1005 library: https://www.ncbi.nlm.nih.gov/clone/library/genomic/234/) (ENA accession:HE000001 – HE565349) [*20*] from the same individual (i.e. Duroc 2-14) were mapped to the contigs and regions with multiple occurrences of incorrect orientation were examined manually in the Integrative Genomics Viewer (IGV) [*61*]. For 28 contigs where there was consistent evidence of structural disagreement between the contigs and the Illumina reads, BAC ends and fosmids, the contigs were split or trimmed.

##### 1.1.3 Scaffolding

In order to establish an initial scaffold the contigs were mapped to Sscrofa10.2 using Nucmer (v3.23) [*27*]. The positioning of the contigs was determined by using the longest ascending subset of mapping locations using the show-coords tool from Mummer with the –g flag. Contigs with a %IDY below 95% were excluded. Contigs that mapped to regions substantially larger (>180%) or smaller (<10%) than the contig size were excluded. These tolerances were intentionally lenient due to the inflated gap sizes in the Sscrofa10.2 assembly (e.g. including 50 kb between scaffolds as required by the NCBI submission system in 2011) and highly fragmented nature of certain regions of Sscrofa10.2. Adjacent contigs were merged into a single fasta entry with Ns representing gaps between them. Gaps were estimated from the distance between the mapping locations against Sscrofa10.2, with an upper limit of 50 kb. Several of the remaining contigs were placed by identifying their longest alignment position, if this alignment was more than 50% the length of the contig and overlapped with a gap with a IDY>90% they were placed in the gaps with 25 bp gaps either side. 346 contigs covering 2.3 Gb were included in the initial chromosomal scaffolds.

##### 1.1.4 Gap filling

PBJelly [*28*] was used with the 65X raw PacBio reads to fill the gaps in the scaffolds. Default parameters were used for all stages except the assembly stage where max wiggle (-w) was set to 100 kb and max trim (-t) was set to 1,000 bp. These parameters were changed to account for the extremely inaccurate gap sizes and missing sequence in Sscrofa10.2 that will have influenced the estimated gap sizes, to allow heavily overlapping contigs to be closed and to allow potentially low-quality sequence at the end of contigs to be excluded. Following initial gap filling, PBJelly was rerun on the fasta output from the first round, with the unused contigs from the Falcon output added to the fasta to allow extension of the scaffolds. These contigs had been excluded initially to reduce secondary mapping positions. PBJelly is able to add contigs to the end of scaffolds, but not place whole contigs in gaps, so the initial mapping of contigs to scaffolds was examined to find if any of the contigs that had been excluded in this stage due to overlap with existing contigs might fill the gaps. Contigs were placed on a case-by-case basis if there was evidence of overlap with placed sequence on both sides of the gap, if the initial contig quality control was good, and if placement was well supported by BAC end mapping. Additionally, BACs for which the end sequences mapped to adjacent contigs providing evidence for scaffolding these adjacent contigs and for which finished quality sequence was publically available, were aligned and the gap filled and placed following the same restrictions as the unplaced contigs. On completion of these gap-filling procedures 108 gaps remained. Estimation of the size of the remaining gaps was based on BAC end mapping, using the known median insert size of the CHORI-242 library (see https://bacpacresources.org). Any gaps estimated to be <100 bp were sized at 100 bp and unspanned gaps were sized at 50 kb.

###### 1.1.5 Targeted BAC sequencing to fill gaps

Five BACs from the CHORI-242 library were selected for further sequencing (CH242-188M9 (SSC16); CH242-323K10 (SSC18); CH242-284F8 (SSC18); CH242-61K12 (SSC1); CH242-168C15 (SSC12)) based on BAC ends mapping either side of gaps. The BAC clones were obtained from BACPAC (https://bacpacresources.org) and DNA was extracted using the Epicentre BACMAX DNA purification kit following manufacturer's instructions. The BAC DNA was sequenced using Oxford Nanopore Technologies' MinION sequencer using a barcoded 2D library following the discontinued protocol SQK-LSK208 on an R9 flow cell using MinKNOW v1.0.5. Sequences were assembled using Canu [*62*] with default settings and each produced a single contig. The BAC vector sequences were removed from the contigs, the contigs were mapped to the assembly initially with Nucmer to confirm they mapped to the expected locations, with exact positions for placement determined by BWA-MEM [*63*]. All five contigs mapped to the expected positions and were placed to close the targeted gaps, leaving 103 gaps in the final Sscrofa11 assembly and closing chromosomes 16 and 18.

###### 1.1.6 Polishing

Error correction was done using Arrow from the GenomicConsensus suite (https://github.com/PacificBiosciences/GenomicConsensus) using the original 65X PacBio coverage. This was followed by Pilon [*64*] with fixlist restricted to "bases", but otherwise using default parameters and paired-end Illumina short read data that provided 50x genome coverage.

#### 1.2 From MARC1423004 DNA to assembly USMARCv1.0

###### 1.2.1 Sample, sequencing and assembly

DNA was isolated from barrow MARC1423004 using a salt extraction method. Briefly, frozen lung tissue was crushed into powder, scraped into a 15 mL tube, and suspended in 4 mL digestion buffer (10 mM $NH_4Cl$, 400 mM NaCl, 50 mM $Na_2EDTA$, pH 8.0). Digestion was initiated with 100 µL 20% SDS and 70 µL trypsin (5 mg/ml). This initial digestion was allowed to proceed at room temperature (approximately 22ºC) for one hour, and then 200 µL of 20% SDS and 50 µL of Proteinase K (50 mg/mL) were added. The digestion was incubated at 55ºC in a shaking water bath overnight (16 hours). Another 100 µL of Proteinase K were added and incubation extended for another 1.5 hours, until no remaining tissue pieces could be observed in the solution, and then 10 µL of RNase (10 U/µL) were added followed by additional incubation for one hour. 1.25 mL 5M NaCl was added, mixed by inversion, and the tube was centrifuged at 3200 x g at 4ºC. The supernatant was transferred to a fresh 15 mL tube, and DNA precipitated by addition of 2.5 volumes of 95% ethanol. The precipitate was removed using a hooked Pasteur pipet, dipped twice in separate tubes of 70% ethanol on ice, and allowed to briefly dry in air on the hook. The DNA was then eluted from the hook by placing it under 250 µL TE buffer (10 mM Tris-HCl, 0.1 mM EDTA) until the pellet slipped off into the buffer. The hook was then removed, and the DNA was allowed to dissolve into the buffer for several days at 4ºC until it appeared to be completely dissolved. The high molecular weight DNA from this extraction was sequenced by Pacific Biosciences (PacBio) using their long read sequencing technology. Libraries for SMRT sequencing were prepared and sequenced as described previously [*48*] using P5/C3 and P6-C4 chemistry on the RSII.  A total of 199 P5/C3 cells and 127 P6/C4 cells were produced. Initial read statistics are detailed in Table S1. Contigs were assembled using Celera Assembler v8.3rc2 [*30*] using the command:

```
wgs-8.3/Linux-amd64/bin/PBcR -s pacbio.spec -fastq
filtered_subreads.fastq genomeSize=3000000000 -sensitive -l swine
sgeName=swine "sge=-p -500 -A swinenewsens"  useGrid=1 scriptOnGrid=1

and spec file:
merSize = 16

ovlMemory = 32
ovlStoreMemory = 32000
ovlThreads = 32
threads = 32
ovlConcurrency = 1
cnsConcurrency = 8
merylThreads = 32
merylMemory = 32000
frgCorrThreads = 16
frgCorrBatchSize = 100000
ovlCorrBatchSize = 100000

useGrid=1
scriptOnGrid=1
ovlCorrOnGrid=1
frgCorrOnGrid=1

sge = -A assembly
sgeScript = -pe threads 1
sgeConsensus = -pe threads 8
sgeOverlap = -pe threads 4 -l mem=2GB
gridEngineMhap = -pe threads 15 -l mem=2GB
sgeCorrection = -pe threads 15 -l mem=2GB
sgeOverlapCorrection  = -pe threads 1 -l mem=16GB
sgeFragmentCorrection=-pe threads 2 -l mem=2GB
sgeOverlapCorrection=-pe threads 1 -l mem=4GB

asmOvlErrorRate=0.1
asmUtgErrorRate=0.06
asmCgwErrorRate=0.1
asmCnsErrorRate=0.1
asmOBT=1
asmObtErrorRate=0.08
asmObtErrorLimit=4.5

batOptions=-RS -NS -CS
utgGraphErrorRate=0.055
utgGraphErrorLimit=4
utgMergeErrorRate=0.055
utgGraphErrorLimit=4

ovlHashBits=24
ovlHashLoad=0.80

ovlHashBlockLength      =300000000
ovlRefBlockLength       =0
```

```
ovlRefBlockSize        =2000000
```

This initial assembly was 2.67 Gbp in 16,441 contigs and an N50 of 2.8 Mbp. Quiver from SMRTportal v. 2.3.0 [*49*] was used to correct the assembly.

##### 1.2.3 Scaffolding

The lung tissue from the pig was sent to Dovetail Genomics (Santa Cruz) for scaffolding by Chicago and HiRise as described [*25*]. This process identified 270 putative misjoins in the contigs and output scaffolds 13,039 scaffolds (294 > 50 kb). The total length was 2.66 Gbp and scaffold N50 was 36.5 Mbp. The dovetail scaffolds were gap-filled where a single contig spanned the gap, correcting false breaks made by HiRise. The resulting assembly was used for reference-guided scaffolding based on the Sscrofa11.1 reference. In case of conflicts, with the exception of cross-chromosome joins, the USDA assembly was unchanged.

##### 1.1.4 Gap filling

PBJelly [*28*] was used with the 65X raw PacBio reads to fill the gaps in the scaffolds. Default parameters were used for all steps.

##### 1.2.5 Polishing

Gap filling was followed by Pilon [*64*] with fixlist restricted to “bases”, but otherwise using default parameters and paired-end Illumina short read data that provided 50x genome coverage. The final assembly of 2.8 Gbp has a scaffold N50 of 131.5 Mbp and a contig N50 of 6.4 Mbp (Table 1).

#### 1.3 Anchoring the assemblies to chromosomes

##### 1.3.1 Chromosome Preparation

Heparinized blood samples were cultured for 72 h in PB MAX Karyotyping medium (Invitrogen) at 37°C, 5% $CO_2$. Cell division was arrested by adding colcemid at a concentration of 10.0 µg/ml (Gibco) for 30 min prior to hypotonic treatment with 75 mM KCl and fixation to glass slides using 3:1 methanol:acetic acid.

##### 1.3.2 Preparation and Selection of BAC clones for FISH

BAC clones with inserts of approximately 150 kb in size were selected for position using the Sscrofa10.2 NCBI database (www.ncbi.nim.nih.gov) and ordered from the PigE-BAC library (ARK-Genomics) [*65*] and the CHORI-242 Porcine BAC library (BACPAC, https://bacpacresources.org/). BAC clone DNA was isolated using the Qiagen Miniprep Kit (Qiagen) prior to amplification and direct labelling by nick translation. Probes were labelled with Texas Red-12-dUTP (Invitrogen) and FITC- Fluorescein-12-UTP (Roche) prior to purification using the Qiagen Nucleotide Removal Kit (Qiagen).

##### 1.3.3 Fluorescence *in situ* hybridisation

Metaphase preparations were fixed to slides and dehydrated through an ethanol series (2 min each in 2×SSC, 70%, 85% and 100% ethanol at RT). Probes were diluted in a formamide buffer (Cytocell) with Porcine Hybloc (Insight Biotech) and applied to the metaphase preparations on a 37°C hotplate before sealing with rubber cement. Probe and target DNA were simultaneously denatured for 2 mins on a 75°C hotplate prior to hybridisation in a humidified chamber at 37°C for 16 h. Slides were washed post hybridisation in 0.4x SSC at 72°C for 2 mins followed by 2x SSC/0.05% Tween 20 at RT for 30 secs, and then counterstained using VECTASHIELD anti-fade medium with DAPI (Vector Labs). Images were captured using an Olympus BX61 epifluorescence microscope with cooled CCD camera and SmartCapture (Digital Scientific UK) system.

### 2. Annotation (Ensembl)

#### 2.1 Repeat Finding

After loading into a database, the pig genome assemblies, including the Sscrofa11.1 reference genome sequence were screened for sequence patterns, including repeats using RepeatMasker (http://www.repeatmasker.org) (version 4.0.5) with parameters '-nolow –species "sus_scrofa" –engine "crossmatch"', Dust (Kuzio, J., R. Tatusov, and D. Lipman, *Dust.* Unpublished but briefly described in [*66*]) and TRF [*52*]. For the pig genome annotation, the Repbase pig library was used with Repeatmasker.

#### 2.2 Low complexity features, ab initio predictions and BLAST analyses

Transcription start sites (TSS) were predicted using Eponine-scan [*67*]. CpG islands [Micklem, G., unpublished] longer than 400 bases and tRNAs [*68*] were also predicted. The results of Eponine-scan, CpG and tRNAscan are for display purposes only and are not used subsequently in the gene annotation process. Genscan [*69*] was run across the repeat-masked sequence to identify *ab initio* gene predictions. The Genscan predictions were used as a supplementary data track, but not used further in the main annotation.

#### 2.3 Protein-coding model generation

Various sources of transcript and protein data were investigated and used to generate gene models using a variety of techniques and are outlined below. Potential transcript models from each evidence source were clustered and each locus was analysed to select the most likely set of full length isoforms, with preference given to species-specific data over data based on homology (described in more detail in section 2.3.9).

##### 2.3.1 cDNA alignment

cDNAs were downloaded from ENA (www.ebi.ac.uk/ena) and RefSeq [*70*], and aligned to the genome using Exonerate [*71*]. Only known mRNAs were used (NMs). A minimal sequence length of 60 bp was used and a cut-off of 97% identity and 90% coverage were required for an alignment to be processed further. Open reading frame potential was assessed via a BLAST of all UniProt protein existence level 1 and 2 vertebrate proteins.

##### 2.3.2 Projection mapping pipeline

Whole genome alignments against the human GRCh38.p12 genome assembly were generated using LastZ. Regions of conserved synteny identified using these alignments were used to map protein-coding annotation from the Ensembl/GENCODE [*72*] gene set. For each protein-coding gene in the human genome, we projected the coding exons within the canonical transcript to pig. In case of exonic overlap on the projected sequence, the longest exon took precedence. Where the mapping did not succeed, we selected the next successful projection of the transcript having the longest translation.

##### 2.3.3 Iso-Seq aligment

PacBio Iso-Seq data are high coverage long read transcriptomic data that allows for correction for the high error rate in raw PacBio reads. The consensus sequences representing nine tissues (brain, diaphragm, hypothalamus, liver, skeletal muscle (*longissimus dorsi*), pituitary, small intestine, spleen, and thymus were downloaded from the short read archive (SRA: PRJNA351265) after correction using Illumina short reads from the same tissue type. The sequences were aligned to the genome using Minimap2 [*73*] with the -splice:hq parameter. Mappings were subsequently filtered to remove sequences with < 90 percent identity and < 95 percent coverage to the reference sequence. All the Iso-Seq data sets had 3' capping and were used for adding UTRs to homology-based protein coding models. All Iso-Seq data sets were used as lncRNA candidates for our lncRNA prediction pipeline.

##### 2.3.4 Protein-to-genome alignment

Protein sequences were downloaded from UniProt and aligned to the genome in a splice aware manner using GenBlast [*74*]. The set of proteins aligned to the genome was a subset of UniProt proteins used to provide a broad targeted coverage of the pig genome. The set consisted of the following:

- Pig SwissProt/TrEMBL PE level 1, 2, 3
- Human SwissProt/TrEMBL PE level 1, 2
- Mouse SwissProt/TrEMBL PE level 1, 2
- Other mammals SwissProt/TrEMBL PE level 1, 2

Note: PE level = protein existence level. A cut-off of 50 percent coverage and identity and an e-value of e-1 were used for GenBlast [*74*] with the exon repair option turned on. The top 5 transcript models built by GenBlast for each protein passing the cut-offs were kept.

##### 2.3.5 RNA-seq alignment

RNA-Seq data were downloaded from ENA (https://www.ebi.ac.uk/ena/) and used in the annotation. RNA-Seq data from PRJEB19386 and PRJEB33353 were used for annotation. The PRJEB19386 RNA-Seq data consisted of 150 bp paired end reads from libraries prepared using a stranded library protocol from ribo-depleted total RNA from Duroc pigs. The PRJEB19386 dataset comprised RNA-Seq data from 28 tissue and cell samples: alveolar macrophages, amygdala, brain stem, caecum, cerebellum, colon, corpus callosum, duodenum, epididymis, frontal lobe (brain), hippocampus, ileum, kidney cortex, left ventricle (heart), mesenteric lymph node, medulla oblongata, occipital lobe, omentum, penis, pituitary gland, pons, skeletal muscle, spleen, stomach, thalamus, tonsil, uterus (Table S9). The PRJEB33353 data set consisted of four tissues brain, muscle, liver and testes. A merged file containing reads from all tissues was also created. The merged data were less likely to suffer from model fragmentation due to read depth. The available reads were aligned to the genome using BWA. A 50 percent allowed mismatch criteria was applied to identify potential splice junctions. Initial rough exon/intron boundaries were generated via the BWA alignments and then refined by mapping the reads in a splice-aware manner using Exonerate [*71*]. Protein-coding models were identified from BLAST alignments of the longest ORF against the UniProt vertebrate PE1 & 2 dataset. Models with poorly scoring or no BLAST alignments were split into a separate class and considered as potential lncRNAs.

##### 2.3.6 Immunoglobulin and T-cell receptor genes

Translations of different human IG gene segments were downloaded from the IMGT database [*75*] and aligned to the genome using GenBlast [*74*]. For the pig annotation, logical thresholds for coverage (80%), percent identity (70%) and e-value (E-1) were used for GenBlast with the exon repair option on. The top 10 transcript models built by GenBlast for each protein passing the cut-offs were kept.

##### 2.3.7 Selenocysteine proteins

Known selenocysteine proteins were aligned against the genome using Exonerate [*71*]. The models generated were checked for the presence of selenocysteines in the same positions as the known proteins. We only kept models with at least 90% coverage and 95% identity.

##### 2.3.8 Filtering the models

The filtering phase decided the subset of protein-coding transcript models, generated from the model-building pipelines, that would comprise the final protein-coding gene set in the GeneBuild. Models were filtered based on information such as what pipeline they were generated using, how closely related the data are to the target species (i.e. pig) and how good the alignment coverage and percent identity to the original data are. Models were filtered using the LayerAnnotation and GeneBuilder modules. The Apollo software [*76*] was used to visualise the results of the filtering.

##### 2.3.9 Collapsing the transcript set

The LayerAnnotation module was used to define a hierarchy of input data sets, from most preferred to least preferred. The output of this pipeline included all transcript models form the highest ranked input set. Models from lower ranked input sets were included only if their exons do not overlap a model from an input set higher in the hierarchy. Note that models cannot exist in more than one layer. For UniProt proteins, models were also separated into clades, to help selection during the layering process.

Each locus was examined in terms of total evidence. The highest priority was given to specialised gene types, the IG/TR genes and selenocysteine genes. After this, priority was given to unique transcript isoforms derived from transcriptomic data (RNA-seq and Iso-Seq) and cDNA sequences that had an open reading frame with a high level of coverage (90 percent or more) to a PE12 vertebrate protein. Gap filling was carried out using homology data from both the human annotation mappings and the UniProt alignments (again with preference for models with 90 percent coverage when re-aligned to the original evidence). After this, potential fragments were added in, again with preference given to the transcriptomic data and cDNAs over the homology data.

A detailed breakdown of the layers used can be found here: https://github.com/Ensembl/ensembl-analysis/blob/feature/pig_98_layers/modules/Bio/EnsEMBL/Analysis/Hive/Config/LayerAnnotationStatic.pm

##### 2.3.10 Addition of UTR to coding models

Models built from homology data lacked untranslated regions (UTRs). In order to call potential UTRs for these cases we used the available cDNA, long and short read data. When multiple UTR donors were available at a locus, the source of the UTR was prioritised with UTR coming from cDNAs and Iso-Seq, then RNA-seq, with a minimum requirement the terminal intron of any spliced models lacking UTR needed to be identical to an intron in the potential UTR donor. If multiple alternative UTRs were available for the highest priority source, suitability of a UTR match was based on the splicing pattern of the UTR donor versus the splicing pattern of the target mode. The donor with the highest number identical splice junctions to the target was selected. If the acceptor ORF was originally missing a start or a stop codon, the ORF was examined for possible extension via the added UTR sequence (while maintaining the frame of the ORF). Single exon ORFs had UTR addition based on residing fully within a potential UTR donor sequence.

##### 2.3.11 Generating multi-transcript genes

The steps described above generated a large set of potential transcript models, many of which overlapped one another. Redundant transcript models were collapsed and the remaining unique set of transcript models were clustered into multi-transcript genes where each transcript in a gene has at least one coding exon that overlaps a coding exon from another transcript within the same genes.

##### 2.3.12 Pseudogenes

Pseudogenes were annotated by looking for genes with evidence of frame-shifting or genes located in regions rich in repetitive sequences. Introns of less than 75 bp length were flagged as potential artificial introns. Transcripts with 2 or more suspect introns or transcripts where >= 80 percent of the introns were covered by repeats where flagged as evidence of a potential pseudogene. Single exon retrotransposed pseudogenes were identified by searching for a multi-exon equivalent elsewhere in the genome.

##### 2.3.13 Small ncRNAs

Small non-coding (sncRNA) genes were added using annotations taken from RFAM [*77*] and miRBase [*78*]. BLAST was run for these sequences to identify homologs in the genome sequence and models were evaluated for expected stem-loop structures using RNAfold [*78*]. Additional machine learning based filters were applied to exclude predictions with sub-optimal alignments to the genome and non-conforming secondary structures. For other sncRNAs, models were built using the Infernal software suite [*80*].

**2.3.14 lncRNAs identification**

Using the transcriptomic data set, we tried to predict long intergenic non-coding RNAs (lncRNAs). We used the RNA-seq and Iso-Seq data which were filtered against the protein-coding gene set. Firstly, candidates with any overlap to a protein alignment evidence used to build the protein coding set were removed. Next sequences that were less than 200bp were removed. Due to the amount of transcriptional noise present in the Iso-Seq a final level of filtration was carried out. Here candidates with less than two splice junctions were removed, unless they had covered at least 1000bp of aligned sequence.

**2.3.15 Cross-referencing and stable identifiers**

Before public release the transcripts and translations were given external references (cross-references to external databases). Stable identifiers were assigned to each gene, transcript, exon and translation. As earlier pig genome sequences have been annotated by Ensembl previously a comparison was made to the previous gene set and as many stable identifiers as possible were mapped between the two annotations.

**2.3.16 Gene expression**

The Illumina RNA-Seq data (Table S9) were also processed by the EBI Gene Expression Atlas (GXA) team [*81*] (https://www.ebi.ac.uk/gxa/home) to generate a baseline gene expression atlas (Expression Atlas release 25, August 2017). These gene expression data can be visualised in the Ensembl genome browser from the gene page.

**2.3.17 Comparison of Ensembl and NCBI annotation**

The Sscrofa11.1 assembly was also annotated independently by the NCBI (https://www.ncbi.nlm.nih.gov/genome/annotation_euk/Sus_scrofa/106/). We have compared these two annotations (Table S7).

#### 2.4 Annotation of non-reference pig assemblies

**2.4.1 Annotation of 11 non-reference pig breed assemblies**

Annotation for 11 other non-reference pig breed assemblies was also carried out using the Ensembl pipeline [*36*]. The breeds included: Bamei, Berkshire, Hampshire, Jinhua, Landrace, Large White, Meishan, Pietrain, Rongchang, Tibetan and Wuzhishan (minipig) (Table S5b). Annotation was carried out using the same code as the reference annotation, using the steps described in the above paragraphs. The only notable differences were: the 'mammals' library was used for repeatmasking and the four tissue PRJEB33353 RNA-seq data set was not used in the breed annotations. These differences are not expected to have had any noticeable effect on the overall result as the tissues in the PRJEB33353 data were already well represented in the other transcriptomic data used. Access to the breed annotations can be found at http://www.ensembl.org/Sus_scrofa/Info/Strains?db=core .

**2.4.2 Annotation of the USMARCv1.0 assembly**

Annotation for USMARCv1.0 was carried out using the Ensembl pipeline *[36]* and the same key steps as outlined for Sscrofa11.1. As the annotation of USMARCv1.0 was carried out prior to the latest reference annotation, the underlying code used had minor differences. These included the use of the 'mammals' library for repeatmasking, using Exonerate instead of Minimap2 for the Iso-Seq alignment and the four tissue PRJEB33353 RNA-seq data set was not used for the USMARCv1.0 annotation. These changes are not expected to have resulted in major differences in terms of the overall results.

#### Supplementary Tables and Figures

**Table S1.** Pacific Biosciences read statistics

| | TJTabasco (Duroc 2-14) | MARC1423004 |
|---|---|---|
| **Chemistry** | P6/C4 | P5/C3 and P6/C4 |
| **Number of reads** | 12,328,735 | 32,960,338 |
| **Total length of reads (bp)** | 175,934,815,397 | 186,973,885,772 |
| **Mean read length (bp)** | 14,270 | 6,144 |
| **Read N50 (bp)** | 19,786 | 9,277 |

**Table S2. Predicted telomeres.** Predicted telomere locations in the Sscrofa11.1 assembly. Number of exact matches of the vertebrate TTAGGG repeat sequence was used to identify candidate telomeres.

| Chr | Start | End | Number of hexamers | Region length (kb) | Strand | Hexamer content |
|---|---|---|---|---|---|---|
| 2 | 151,924,806 | 151,935,981 | 1,609 | 11.2 | + | 86.4% |
| 3 | 132,840,959 | 132,848,913 | 1,046 | 8.0 | + | 78.9% |
| 6 | 170,835,933 | 170,843,587 | 957 | 7.7 | + | 75.0% |
| 8 | 138,963,948 | 138,966,197 | 208 | 2.2 | + | 55.5% |
| 9 | 139,499,115 | 139,512,083 | 1,836 | 13.0 | + | 84.9% |
| 14 | 141,745,369 | 141,755,446 | 1,201 | 10.1 | + | 71.5% |
| 15 | 140,408,314 | 140,412,713 | 595 | 4.4 | + | 81.2% |
| 18 | 55,971,782 | 55,982,971 | 1,571 | 11.2 | + | 84.2% |
| X | 125,929,106 | 125,939,592 | 1,329 | 10.5 | + | 76.0% |

**Table S3. Predicted centromeres.** Predicted centromere locations in the Sscrofa11.1 assembly.

| Chr | Start | End | Repeat content (bp) | Region length (bp) | Repeat content |
|---|---|---|---|---|---|
| 1 | 92,615,481 | 92,672,216 | 46,164 | 56,735 | 81.4% |
| 1 | 92,760,768 | 92,881,119 | 110,990 | 120,351 | 92.2% |
| 1 | 93,266,464 | 93,430,514 | 80,940 | 16,4050 | 49.3% |
| 2 | 50,550,173 | 50,777,308 | 198,336 | 227,135 | 87.3% |
| 3 | 41,776,737 | 41,860,603 | 35,376 | 83,866 | 42.2% |
| 4 | 46,443,460 | 46,472,085 | 28,625 | 28,625 | 100.0% |
| 5 | 39,774,025 | 39,828,563 | 54,538 | 54,538 | 100.0% |
| 5 | 39,878,566 | 40,207,105 | 328,539 | 328,539 | 100.0% |
| 6 | 38,712,705 | 38,886,534 | 163,335 | 173,829 | 94.0% |
| 7 | 24,578,125 | 24,606,761 | 28,636 | 28,636 | 100.0% |
| 8 | 144 | 20,905 | 20,761 | 20,761 | 100.0% |
| 8 | 54,585,508 | 54,685,241 | 21,099 | 99,733 | 21.2% |
| 9 | 63,144,551 | 63,503,859 | 356,770 | 359,308 | 99.3% |
| 11 | 11,220,831 | 11,222,126 | 1,295 | 1,295 | 100.0% |
| 11 | 35,726,738 | 35,728,355 | 1,617 | 1,617 | 100.0% |
| 11 | 35,804,210 | 35,809,503 | 5,293 | 5,293 | 100.0% |
| 11 | 35,870,705 | 35,878,206 | 7,501 | 7,501 | 100.0% |
| 13 | 34 | 152,474 | 150,375 | 152,440 | 98.6% |
| 15 | 1,649 | 36,105 | 10,369 | 34,456 | 30.1% |
| 15 | 56,407,100 | 56,427,869 | 9,798 | 20,769 | 47.2% |
| 17 | 63,189,675 | 63,361,433 | 171,758 | 171,758 | 100.0% |
| 18 | 619 | 17,212 | 16,593 | 16,593 | 100.0% |
| Y | 42,496,777 | 42,515,903 | 17,954 | 19,126 | 93.9% |

**Table S4. Assigning scaffolds to chromosomes.** Fluorescent *in situ* hybridisation results using named BAC clones as probes plus sequence matches for sequences derived from these BAC clones.

| Chr | BAC Name | BES | FISH | Sscrofa11.1 coordinates | USMARCv1.0 coordinates |
|---|---|---|---|---|---|
| 1 | PigE-232G23 | CT070230.1; CT218278.1 | 1p | 1:615,021-619,597 | 1:280,453,704-280,458,272 |
| 1 | CH242-248F13 | FP340244.3 | 1p | 1:1,470,202-1,660,001 | 1:279,368,385-279,558,294 |
| 1 | CH242-151E10 | CT239299.1; CT245986.1 | 1q | unplaced scaffold: Contig1206 | 1: 6,156,768-6,336,737 |
| 2 | PigE-117G14 | CT074446.1; CT074447.1 | 2p | 2:19,406-161,226 | 2:537,026-678,808 |
| 2 | PigE-8G19 | CT260033.1; CT260032.1 | 2p | 2:552,031-671,098 | 2:29,620-146,529 |
| 2 | CH242-188K23 | CU929880 | 2 cen | 2:52,747,463-52,933,130 | 2:51,728,649-51,908,148 |
| 2 | CH242-230M23 | CT144824.1; CT258059.1 | 2 cen | 2:53,300,582-53,472,497 | no match |
| 2 | CH242-441A1 | CT364255.1;CT364256.1 | 2 cen | 2:53,458,574-53,652,606 | 2:52,095,251-52,095,932 |
| 2 | CH242-294F6 | CT378635.1; CT378634.1 | 2q | 2:151,178,736-151,402,963 | 2:145,456,152-145,678,427 |
| 3 | PigE-168G22 | CT094069.1; CT094070.1 | 3p | 3:301,813-509,346 | 3:218,358-425,025 |
| 3 | CH242-315N8 | CT359002.1; CT359003.1 | 3q | 3:122,720,374-122,869,530 | no match |
| 4 | PigE-262E12 | CT082779.1; CT193441.1 | 4p | 4:37,383-223,717 | 4: 96,811- 97,511 |
| 4 | PigE-131J18 | CT116562.1; CT171811.1 | 4p | 4:449,934-626,677 | 4:322,853-499,367 |
| 4 | PigE-85G21 | CT070098.1; CT190031.1 | 4q | 4:130,625,653-130,748,215 | 4:130,215,908-130,338,404 |
| 5 | CH242-288F8 | CT132004.1; CT211915.1 | 5p | 5:170,319-344,353 | 5:188,019-362,653 |
| 5 | PigE-178M22 | CT139068.1; CT155898.1 | 5p | 5:175,168-311,462 | 5:192,886-329113 |
| 5 | CH242-133F9 | CT166002.1; CT166003.1 | 5p | 5:438,296-633,458 | 5:456,924-652,458 |
| 5 | PigE-127K14 | CT057696.1; CT057697.1 | 5p | 5:1,003,455-1,129,329 | 5:1,024,261-1,148,699 |
| 5 | PigE-74P10 | CT188857.1; CT188858.1 | 5p | 5:3,739,938-3,883,755 | 5:103,338,585-103,481,984 |
| 5 | PigE-99L23 | CT079916.1; CT106700.1 | 5p Mid | 5:31,980,969-32,114,628 | no match |
| 5 | CH242-63B20 | FP102738 | 5q | 5:104,304,289-104,489,770 | no match |
| 6 | PigE-238J17 | CT220438.1; CT220439.1 | 6p | 6:2,333,972-2,522,065 | 6:162,952,836-163,141,204 |
| 6 | PigE-199E24 | CT272854.1; CT272853.1 | 6 below cen | 6:62,771,286-62,952,647 | 6:104,969,580-105,152,317 |
| 6 | CH242-510F2 | CT396711.1; CT442620.1 | 6q | 6:170,248,061-170,454,571 | 6:162,654-369,119 |
| 7 | PigE-52L22 | CT054562.1; CT063652.1 | 7p | 7:188,339-317,255 | 7:125,463,765-125,463,765 |
| 7 | PigE-246A1 | CT203984.1; CT070741.1 | 7 cen | 7:24,628,314-24,671,828 | no match |
| 7 | PigE-230H8 | CT120917.1 | 7q below cen | 7:46,704,415-46,704,995 | 7:395,704-396,284 |
| 7 | PigE-75E21 | CT188956.1; CT261917.1 | 7q below cen | 7:46,901,592-47,032,091 | 7:68,406-199,212 |
| 7 | CH242-103I13 | CU695123.2 | 7q | Unplaced scaffold: Contig1914 | 7:7,614,911-7,838,927 |

| Chr | BAC Name | BES | FISH | Sscrofa11.1 coordinates | USMARCv1.0 coordinates |
|---|---|---|---|---|---|
| 8 | PigE-134L21 | CT126839.1; CT172501.1 | 8p | 8:570,904-705,341 | 8:280,369,080-280,502,409 |
| 8 | PigE-2N1 | CT229915.1; CT229916.1 | 8p | 8:819,717-958,131 | 8:137,599,822-137,737,820 |
| 8 | PigE-118B21 | CT048761.1; CT091504.1 | 8q | 8:138,491,413-138,647,394 | 8:322,914-478,869 |
| 9 | CH242-65G4 | CU695192.2 | 9p | 9:320,582-511,079 | 9:137,686,630-137,874,917 |
| 9 | PigE-126O17 | CT170583.1; CT057320.1 | 9p | 9:443,462-603,022 | 9:137,594,779-137,754,110 |
| 9 | PigE-242D8 | CT123266.1; CT123265.1 | 9 mid | 9:67,752,381-67,910,109 | 9:71,096,887-71,254,731 |
| 9 | CH242-411M8 | CT362997.1; CT468791.1 | 9q | 9:139,180,446-139,338,710 | 9:168,756-327,007 |
| 10 | CH242-451I23 | CT369304.1; CT459538.1 | 10p | Unplaced scaffold: Contig2471 | 10:71,863,534-72,028,842 |
| 10 | CH242-36D16 | CT345373.1; CT186999.1 | 10q | 10:55,422,866-55,600,351 | 10:15,300,371-15,480,359 |
| 10 | CH242-517L16 | FP325295.2 | 10q | 10:55,609,778-55,800,022 | 10:15,098,969-15,290,916 |
| 11 | PigE-199B10 | CT272693.1 | 11p | 11:135,233-297,713 | 11:79,101,520-79,264,254 |
| 11 | PigE-232N19 | CT193346.1 | 11p | 11:290,540-291,222 | 11:79,108,017-79,108,697 |
| 11 | PigE-211E21 | CT044498.1; CT044499.1 | 11p | 11:1,584,043-1,743,425 | 11:77,663,220-77,822.434 |
| 11 | CH242-239O11 | CT146353.1; CT286242.1 | 11q | 11:78,888,491-79,057,526 | 11:827,483-996,382 |
| 12 | PigE-253K5 | CT081057.1; CT204391.1 | 12p | 12:324,614-524,015 | 12:3,288-206,400 |
| 12 | PigE-124G15 | CT056668.1; CT092177.1 | 12q | 12:60,846,540-60,990,610 | 12:58,746,918-58,890,342 |
| 13 | PigE-197C11 | CT271598.1; CT271599.1 | 13p | 13:556,804-694,010 | 13:204,579,401-204,716,338 |
| 13 | PigE-179J15 | CT124924.1; CT124925.1 | 13q | 13:205,856,740-206,006,912 | 13:3,005,553-3,154,893 |
| 14 | PigE-137C12 | FP340551.3 | 14p | 14:17,423-156,591 | 14:140,940,126-140,804,938 |
| 14 | PigE-167E18 | CT089616.1; CT089617.1 | 14q | 14:141,407,495-141,435,234 | 14:98,899-125,652 |
| 15 | PigE-90C11 | CT190903.1; CT190904.1 | 15p | 15:3,442,144-3,596,666 | 15:139,733,189-139,886,921 |
| 15 | PigE-108N22 | CT073138.1; CT046453.1 | 15 mid | 15:56,903,229-57,028,679 | no match |
| 15 | CH242-170N3 | FP236135.2 | 15q | 15:139,616,279-139,784,756 | 15:3,511,408-3,588,855 |
| 16 | PigE-90L22 | CT191132.1; CT113297.1 | 16p | 16:109,696-235,547 | 16:87,402-212,531 |
| 16 | PigE-124C22 | CT056551.1; CT056550.1 | 16p | 16:117,329-308,428 | 16:94,873-287,243 |
| 16 | CH242-4G9 | CT041970.1; CT041969.1 | 16p | 16:141,557-324,802 | 16:118,753-303,587 |
| 16 | PigE-173H6 | CT123878.1; CT123877.1 | 16p | 16:167,106-299,570 | 16:144,276-278,432 |
| 16 | PigE-149F10 | CT088298.1; CT153977.1 | 16p | 16:596,671-782,524 | 16:78,918,129-79,108,868 |
| 16 | CH242-42L16 | CT347302.1; CT347303.1 | 16q | 16:79,097,179-79,303,695 | 16:878,687-1,085,418 |
| 17 | CH242-70L7 | CT077340.1; CT077341.1 | 17p | 17:545,995-673,770 | 17:464,378-592,438 |
| 17 | PigE-190G24 | CT126644.1; CT096362.1 | 17p | 17:515,422-707,787 | 17:433,829-626,496 |
| 17 | CH242-243H19 | CT321876.1; CT321877.1 | 17q | 17:61,760-582-61,937,945 | 17:62,450,941-62,628,249 |

| Chr | BAC Name | BES | FISH | Sscrofa11.1 coordinates | USMARCv1.0 coordinates |
|---|---|---|---|---|---|
| 18 | PigE-253N22 | CT081116.1; CT204433.1 | 18p | 18:1,616,389-1,751,286 | 18:1,565,719-1,700,920 |
| 18 | PigE-202I11 | CT042866.1; CT254626.1 | 18q | 18:55,539,630-55,700,409 | 18:55,320,418-55,481,057 |
| X | CH242-447L20 | CT377508.1; CT467360.1 | Xp | X:505,086-692,549 | no match |
| X | CH242-156O11 | FP074895.7 | Xp + Yp | X:6,337,709 6,584,993 | X:7,588,110-7,597,109 |
| X | CH242-19N1 | CU856094.8 | Xp | X:6,705,194-6,834,183 | X:7,588,110-7,715,932 |
| X | CH242-305A15 | CU861979.13 | Xq | X:125,384,028-125,529,813 | X:126,150,718-126,296,945 |
| Y | CH242-156O11 | FP074895.7 | Xp + Yp | Y:4,744,231-4,791,971 | Y:32,909,634-32,923,401 |

**Table S5a. Assemblytics comparisons.** Index to results of comparisons between Sscrofa11.1 and twelve other assemblies

| **Reference** | | **Sscrofa10.2 (GCF_000003025.5)** |
|---|---|---|
| **Query** | **Assembly accession** | |
| **Sscrofa11.1** | GCA_000003025.6 | http://qb.cshl.edu/assemblytics/analysis.php?code=i0H3KuHhWjKO5Tn7nsXg |
| **USMARCv1.0** | GCA_002844635.1 | http://qb.cshl.edu/assemblytics/analysis.php?code=faROmPzOlMp1q5IdToO8 |

| **Reference** | | **Sscrofa11.1** |
|---|---|---|
| **Query** | **Assembly accession** | |
| **Sscrofa11.1** | GCA_000003025.6 | N/A |
| **USMARCv1.0** | GCA_002844635.1 | http://assemblytics.com/analysis.php?code=4rscWrlT7paorSvTMI7L |
| **Bamei** | GCA_001700235.1 | http://assemblytics.com/analysis.php?code=gpCq8VWG4aWrocrlCWww |
| **Berkshire** | GCA_001700575.1 | http://assemblytics.com/analysis.php?code=dvVxU3qkCNUR3rWpm2FI |
| **Hampshire** | GCA_001700165.1 | http://assemblytics.com/analysis.php?code=V6jWeDYKywLu4Av40Ikh |
| **Jinhua** | GCA_001700295.1 | http://qb.cshl.edu/assemblytics/analysis.php?code=UxtEbFk065DWQBpYz0sV |
| **Landrace** | GCA_001700215.1 | http://qb.cshl.edu/assemblytics/analysis.php?code=7V7QGUCXrNAtFcGL6DMT |
| **LargeWhite** | GCA_001700135.1 | http://qb.cshl.edu/assemblytics/analysis.php?code=UymCHs1NirQkdMFFbM1e |
| **Meishan** | GCA_001700195.1 | http://qb.cshl.edu/assemblytics/analysis.php?code=toDVmO7nus0BbyMCGKSc |
| **Pietrain** | GCA_001700255.1 | http://qb.cshl.edu/assemblytics/analysis.php?code=TIlXYB2uQYgWbf5YqNXk |
| **Rongchang** | GCA_001700155.1 | http://qb.cshl.edu/assemblytics/analysis.php?code=HzggG8kBPJ6uKWWEvZOV |
| **Tibetan** | GCA_000472085.2 | http://qb.cshl.edu/assemblytics/analysis.php?code=o9WtyIF6wTnGsEeAiizn |
| **Wuzhishan** | GCA_000325925.2 | http://qb.cshl.edu/assemblytics/analysis.php?code=UbH3avfeoW19DjJmVC8C |

**Table S5b. Assemblytics comparisons.** Index to results of comparisons between USMARCv1.0 and twelve other assemblies

| **Reference** | | USMARCv1.10 |
|---|---|---|
| **Query** | **Assembly accession** | |
| **Sscrofa11.1** | GCA_000003025.6 | http://assemblytics.com/analysis.php?code=4rscWrIT7paorSvTMI7L |
| **USMARCv1.0** | GCA_002844635.1 | N/A |
| **Bamei** | GCA_001700235.1 | http://assemblytics.com/analysis.php?code=A1doW581DPkQKXIwfbtB |
| **Berkshire** | GCA_001700575.1 | http://qb.cshl.edu/assemblytics/analysis.php?code=5dCXFbth2110zsguw58t |
| **Hampshire** | GCA_001700165.1 | http://qb.cshl.edu/assemblytics/analysis.php?code=Xe5ENqAjsxeNcrK7TaRp |
| **Jinhua** | GCA_001700295.1 | http://qb.cshl.edu/assemblytics/analysis.php?code=nqEihnLJRPsjNswVxV9J |
| **Landrace** | GCA_001700215.1 | http://qb.cshl.edu/assemblytics/analysis.php?code=tfrtkAXiy148TUsb8HIJ |
| **LargeWhite** | GCA_001700135.1 | http://qb.cshl.edu/assemblytics/analysis.php?code=IZM3EFMBzo9KyytQMSWH |
| **Meishan** | GCA_001700195.1 | http://qb.cshl.edu/assemblytics/analysis.php?code=K9qeCrVxr9znPtFanHd3 |
| **Pietrain** | GCA_001700255.1 | http://qb.cshl.edu/assemblytics/analysis.php?code=U1n9D7z7DtRvbWjqEdTH |
| **Rongchang** | GCA_001700155.1 | http://qb.cshl.edu/assemblytics/analysis.php?code=nEk3faE5s8YYckjNuvN7 |
| **Tibetan** | GCA_000472085.2 | http://qb.cshl.edu/assemblytics/analysis.php?code=NqjCZ7wvt6D0vm7Ai4tN |
| **Wuzhishan** | GCA_000325925.2 | http://qb.cshl.edu/assemblytics/analysis.php?code=mEqp9WaGi9eceSY4Vid6 |

**Table S5c. Assembly statistics.** Assembly statistics* for pig genome assemblies subject to Assemblytics analyses

| Assembly | Accession | Total (bp)(ungapped) | Scaffolds | Scaffold N50 | Contigs | Contig N50 |
|---|---|---|---|---|---|---|
| **Sscrofa11.1** | GCA_000003025.6 | 2,472,047,747 | 706 | 88,231,837 | 1,118 | 48,231,277 |
| **USMARCv1.0** | GCA_002844635.1 | 2,623,130,238 | 14,818 | 131,458,098 | 14,818 | 6,372,407 |
| **Bamei** | GCA_001700235.1 | 2,433,636,520 | 129,335 | 1,529,027 | 187,466 | 70,893 |
| **Berkshire** | GCA_001700575.1 | 2,414,739,650 | 94,468 | 1,655,397 | 137,661 | 94,651 |
| **Hampshire** | GCA_001700165.1 | 2,418,011,428 | 82,206 | 1,550,023 | 122,452 | 102,417 |
| **Jinhua** | GCA_001700295.1 | 2,433,032,022 | 115,554 | 1,478,908 | 158,796 | 95,227 |
| **Landrace** | GCA_001700215.1 | 2,420,570,845 | 94,659 | 1,407,841 | 141,909 | 88,142 |
| **LargeWhite** | GCA_001700135.1 | 2,430,896,979 | 102,342 | 2,441,555 | 150,742 | 88,831 |
| **Meishan** | GCA_001700195.1 | 2,438,814,343 | 133,833 | 1,248,180 | 201,146 | 63,263 |
| **Pietrain** | GCA_001700255.1 | 2,415,062,022 | 88,436 | 1,663,542 | 139,497 | 80,611 |
| **Rongchang** | GCA_001700155.1 | 2,429,730,895 | 120,246 | 2,325,000 | 173,508 | 79,093 |
| **Tibetan** | GCA_000472085.2 | 2,379,878,366 | 72,068 | 861,885 | 148,234 | 57,199 |
| **Wuzhishan** | GCA_000325925.2 | 2,453,484,489 | 137,577 | 5,853,977 | 272,163 | 31,939 |

* source NCBI Assembly

**Table S6. BUSCO results.** BUSCO statistics, BUSCOv2 (OrthoDBv9).

| | Pig Sscrofa10.2 | Pig Sscrofa11.1 | Pig USMARCv1.0 | Human GRCh38p5 | Mouse GRCm39p3 |
|---|---|---|---|---|---|
| **Complete BUSCOs** | 80.9% | 93.8% | 93.1% | 94.9% | 95.2% |
| **Complete and single-copy BUSCOs** | 80.2% | 93.3% | 92.6% | 94.1% | 91.6% |
| **Complete and duplicated BUSCOs** | 0.7% | 0.5% | 0.5% | 0.8% | 3.6% |
| **Fragmented BUSCOs** | 8.2% | 3.5% | 3.5% | 2.5% | 2.3% |
| **Missing BUSCOs** | 10.9% | 2.7% | 3.4% | 2.6% | 2.5% |
| **Total BUSCO groups searched** | 4,104 | 4,104 | 4,104 | 4,104 | 4,104 |

**Table S7. Annotation statistics.** Comparison of Ensembl and NCBI annotation of Sscrofa11.1.

| **Ensembl** | | **NCBI** | | | |
|---|---|---|---|---|---|
| | | | missing (relative location) | | |
| | | in common | (intragenic) | (intergenic) | other |
| Protein-coding | 21,301 | 17,676 | 468 | 2,617 | [1]540 |
| Non-coding | 8,971 | 1,700 | 627 | 6,346 | [2]298 |
| Pseudogenes | 1,626 | 134 | 4 | 1,193 | [3]295 |
| **NCBI** | | **Ensembl** | | | |
| | | | missing (relative location) | | |
| | | in common | (intragenic) | (intergenic) | other |
| Protein-coding | 20,790 | 17,676 | 140 | 2,522 | [4]452 |
| Non-coding | 6,460 | 1,700 | 100 | 4,433 | [5]227 |
| Pseudogenes | 3,084 | 134 | 4 | 2,884 | [6]62 |

[1] total of 540 protein-coding genes of which 145 were annotated as non-coding by NCBI and the remaining 395 were annotated as pseudogenes by NCBI

[2] total of 298 non-coding genes of which 271 were annotated as protein-coding by NCBI and the remaining 27 were annotated as pseudogenes by NCBI

[3] total of 295 pseudogenes genes of which 281 were annotated as protein-coding by NCBI and the remaining 14 were annotated as non-coding by NCBI

[4] total of 452 protein-coding genes of which 222 were annotated as non-coding by Ensembl and the remaining 230 were annotated as pseudogenes by Ensembl

[5] total of 227 non-coding genes of which 215 were annotated as protein-coding by Ensembl and the remaining 12 were annotated as pseudogenes by Ensembl

[6] total of 62 pseudogenes genes of which 34 were annotated as protein-coding by Ensembl and the remaining 28 were annotated as non-coding by Ensembl

**Table S8. Commercial SNP chip probes.** SNP chip markers mapped to pig genome assemblies.

| Assembly | Mapped / unmapped | AxiomHD | PorcineSNP60 | GGP LD | 80K |
|---|---|---|---|---|---|
| **Sscrofa10.2** | mapped | 633,705 | 59,590 | 50,530 | 68,046 |
| | unmapped | 24,987 | 1,975 | 385 | 470 |
| **Sscrofa11.1** | mapped | 628,280 | 61,299 | 50,586 | 68,270 |
| | ummapped | 30,412 | 266 | 329 | 246 |
| **USMARCv1.0** | mapped | 618,771 | 60,692 | 50,042 | 67,604 |
| | unmapped | 39,921 | 873 | 873 | 912 |

**Table S9. Tissue samples for PRJEB19386.** Tissue samples characterised by Illumina short read RNA-Seq analyses.

| Tissue | BioSample accession | alias | Animal | Sex |
|---|---|---|---|---|
| alveolar macrophages | SAMEA103886124 | SUS_RI_DUR21-30 | Duroc 21 | female |
| alveolar macrophages | SAMEA103886168 | SUS_RI_Pig 21_DUR_30 | Duroc 21 | female |
| alveolar macrophages | SAMEA103886137 | SUS_RI_DUR22-60 | Duroc 22 | male |
| alveolar macrophages | SAMEA103886112 | SUS_RI_Pig 22_DUR_60 | Duroc 22 | male |
| amygdala | SAMEA103886173 | SUS_RI_R-Dur_23-08 | Duroc 23 | female |
| amygdala | SAMEA103886162 | SUS_RI_Dur_24-C-S0 | Duroc 24 | male |
| brain, frontal lobe | SAMEA103886139 | SUS_RI_R-Dur_23-01 | Duroc 23 | female |
| brain, frontal lobe | SAMEA103886156 | SUS_RI_R-Dur_24-41 | Duroc 24 | male |
| brain stem | SAMEA103886128 | SUS_RI_R-Dur_23-05 | Duroc 23 | female |
| brain stem | SAMEA103886129 | SUS_RI_R-Dur_24-45 | Duroc 24 | male |
| caecum | SAMEA103886133 | SUS_RI_DUR21-19 | Duroc 21 | female |
| caecum | SAMEA103886120 | SUS_RI_DUR22-48 | Duroc 22 | male |
| caecum | SAMEA103886151 | SUS_RI_Pig 22_DUR_48 | Duroc 22 | male |
| cerebellum | SAMEA103886116 | SUS_RI_R-Dur_23-09 | Duroc 23 | female |
| cerebellum | SAMEA103886131 | SUS_RI_R-Dur_24-49 | Duroc 24 | male |
| colon | SAMEA103886132 | SUS_RI_Dur_23-21 | Duroc 23 | female |
| colon | SAMEA103886147 | SUS_RI_Dur_24-61 | Duroc 24 | male |
| corpus callosum | SAMEA103886154 | SUS_RI_R-Dur_23-10 | Duroc 23 | female |
| corpus callosum | SAMEA103886167 | SUS_RI_R-Dur_24-50 | Duroc 24 | male |
| duodenum | SAMEA103886155 | SUS_RI_Dur_23-22 | Duroc 23 | female |
| duodenum | SAMEA103886176 | SUS_RI_Dur_24-62 | Duroc 24 | male |
| epididymis | SAMEA103886140 | SUS_RI_DUR22-58 | Duroc 22 | male |
| hippocampus | SAMEA103886122 | SUS_RI_Dur_23-B-S0 | Duroc 23 | female |
| hippocampus | SAMEA103886114 | SUS_RI_R-Dur_24-51 | Duroc 24 | male |
| ileum | SAMEA103886163 | SUS_RI_Dur_23-23 | Duroc 23 | female |
| ileum | SAMEA103886121 | SUS_RI_Dur_24-63 | Duroc 24 | male |
| kidney cortex | SAMEA103886174 | SUS_RI_DUR21-09 | Duroc 21 | female |
| kidney cortex | SAMEA103886153 | SUS_RI_DUR22-39 | Duroc 22 | male |
| heart, left ventricle | SAMEA103886169 | SUS_RI_DUR21-12 | Duroc 21 | female |
| heart, left ventricle | SAMEA103886172 | SUS_RI_DUR22-43 | Duroc 22 | male |
| lymph node, mesenteric | SAMEA103886127 | SUS_RI_DUR21-22 | Duroc 21 | female |
| lymph node, mesenteric | SAMEA103886115 | SUS_RI_DUR22-51 | Duroc 22 | male |
| medulla oblongata | SAMEA103886135 | SUS_RI_R-Dur_23-06 | Duroc 23 | female |
| medulla oblongata | SAMEA103886142 | SUS_RI_R-Dur_24-46 | Duroc 24 | male |

| Tissue | BioSample accession | alias | Animal | Sex |
|---|---|---|---|---|
| occipital lobe | SAMEA103886158 | SUS_RI_R-Dur_23-02 | Duroc 23 | female |
| occipital lobe | SAMEA103886177 | SUS_RI_R-Dur_24-42 | Duroc 24 | male |
| omentum | SAMEA103886145 | SUS_RI_DUR21-65 | Duroc 21 | female |
| omentum | SAMEA103886146 | SUS_RI_DUR22-73 | Duroc 22 | male |
| penis | SAMEA103886166 | SUS_RI_DUR22-59 | Duroc 22 | male |
| pituitary gland | SAMEA103886152 | SUS_RI_Dur_23-14 | Duroc 23 | female |
| pituitary gland | SAMEA103886150 | SUS_RI_Dur_24-54 | Duroc 24 | male |
| pituitary gland | SAMEA103886149 | SUS_RI_DUR21-06 | Duroc 21 | female |
| pons | SAMEA103886159 | SUS_RI_R-Dur_23-07 | Duroc 23 | female |
| pons | SAMEA103886164 | SUS_RI_R-Dur_24-47 | Duroc 24 | male |
| skeletal muscle | SAMEA103886171 | SUS_RI_DUR21-24 | Duroc 21 | female |
| skeletal muscle | SAMEA103886118 | SUS_RI_DUR22-75 | Duroc 22 | male |
| spleen | SAMEA103886157 | SUS_RI_DUR21-25 | Duroc 21 | female |
| spleen | SAMEA103886170 | SUS_RI_DUR22-55 | Duroc 22 | male |
| stomach | SAMEA103886111 | SUS_RI_Dur_23-24 | Duroc 23 | female |
| stomach | SAMEA103886134 | SUS_RI_Dur_24-64 | Duroc 24 | male |
| thalamus | SAMEA103886136 | SUS_RI_R-Dur_23-13 | Duroc 23 | female |
| thalamus | SAMEA103886160 | SUS_RI_R-Dur_24-53 | Duroc 24 | male |
| tonsils | SAMEA103886125 | SUS_RI_DUR22-56 | Duroc 22 | male |
| uterus | SAMEA103886126 | SUS_RI_DUR21-27 | Duroc 21 | female |

**Figure S1. Predicted telomeres.** Predicted locations of telomeres in the Sscrfoa11.1 assembly

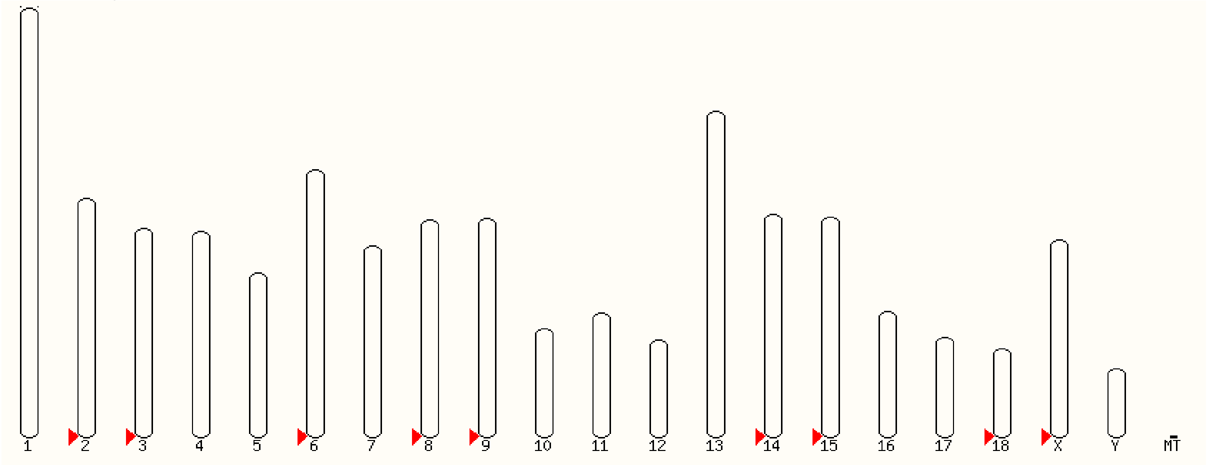

**Figure S2. Predicted centromeres.** Predicted centromere locations in the Sscrofa11.1 assembly.

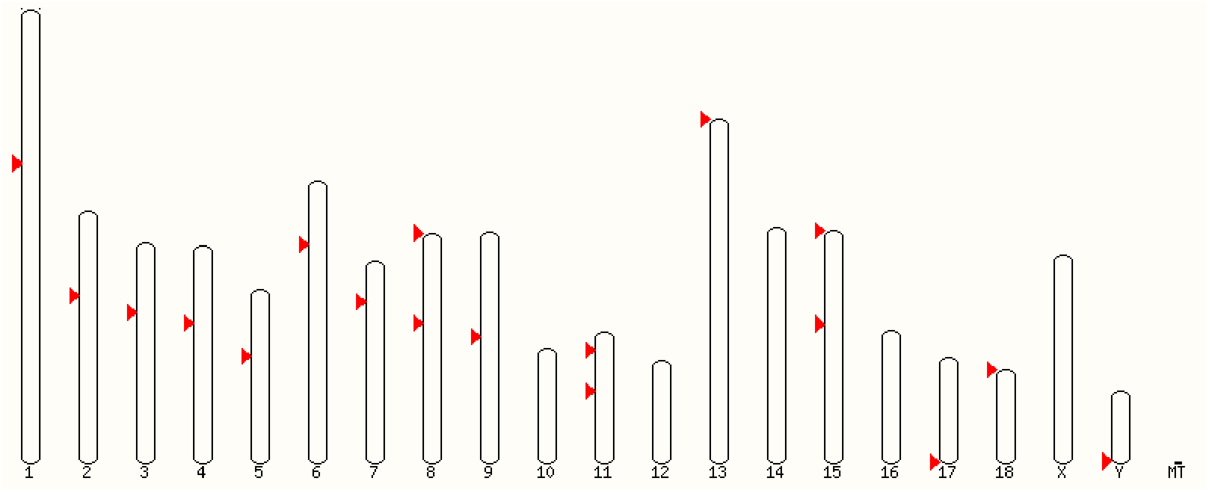

**Figure S3. Fluorescent *in situ* hybridisation assignments.**

**a.** SSC6 – p-telomeric end labelled with PigE-238J17, q-telomeric end labelled with CH242-510F2

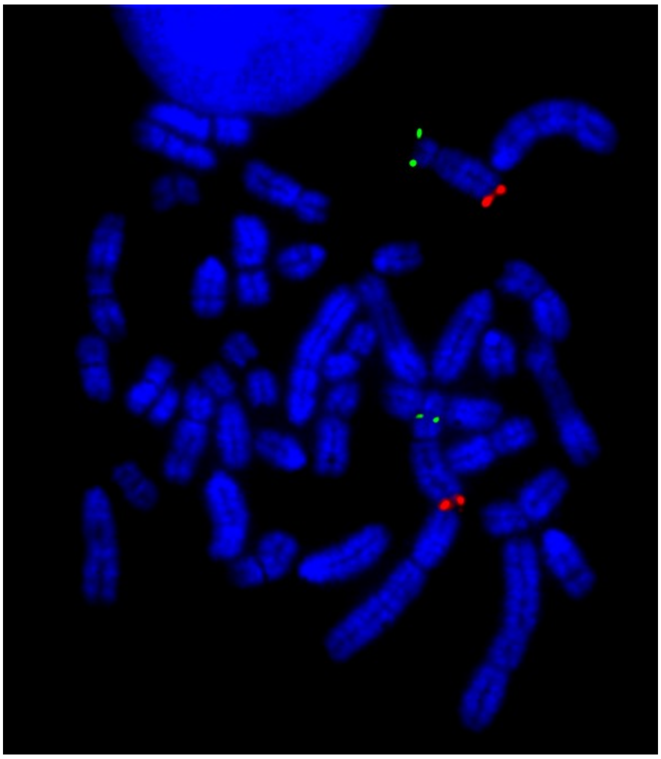

**b.** SSCX – p-telomeric end labelled with CH242-19N1, q-telomeric end labelled with CH242-305A15

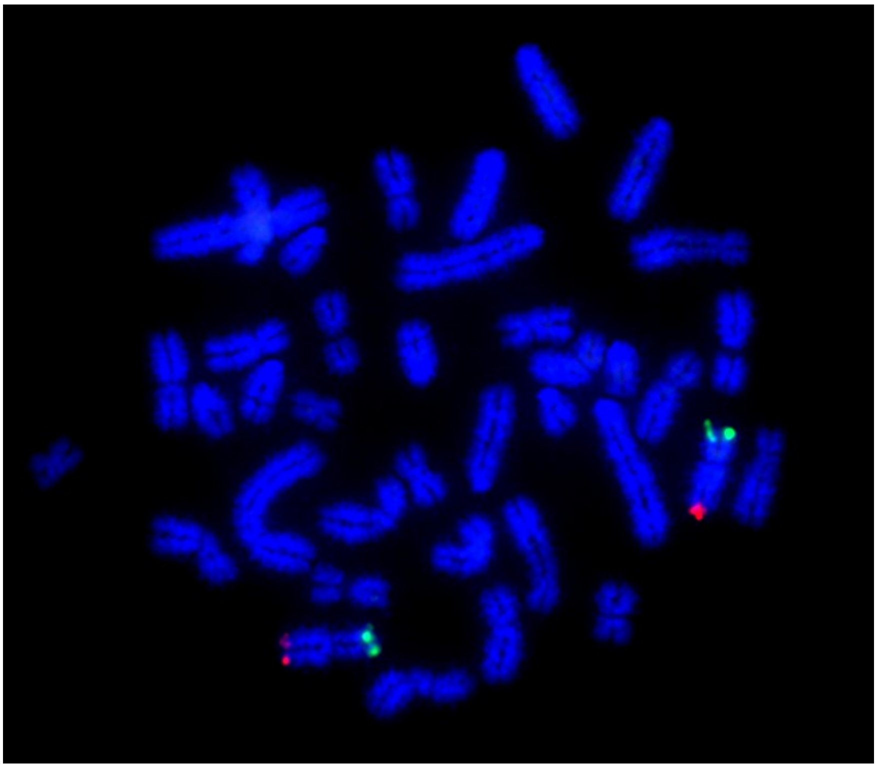

**Fig. S4. Improvement in local order and orientation and reduction in redundancy**. gEVAL screenshort showing the alignment of isogenic CH242 BAC end and WTSI_1005 fosmid end sequences with the Sscrofa10.2 (upper panel with pink bar on left hand side) and Sscrofa11.1 (lower panel). Red arrows indicate incorrect orientation of the paired end sequences, purple arrows are sequences which are present multiple times, green and orange arrows indicate the end sequences are correctly oriented. The distances between correctly oriented end sequences are as expected (green) or either greater or less than expected (orange) for the clone insert size for the fosmid or CH242 BAC libraries.

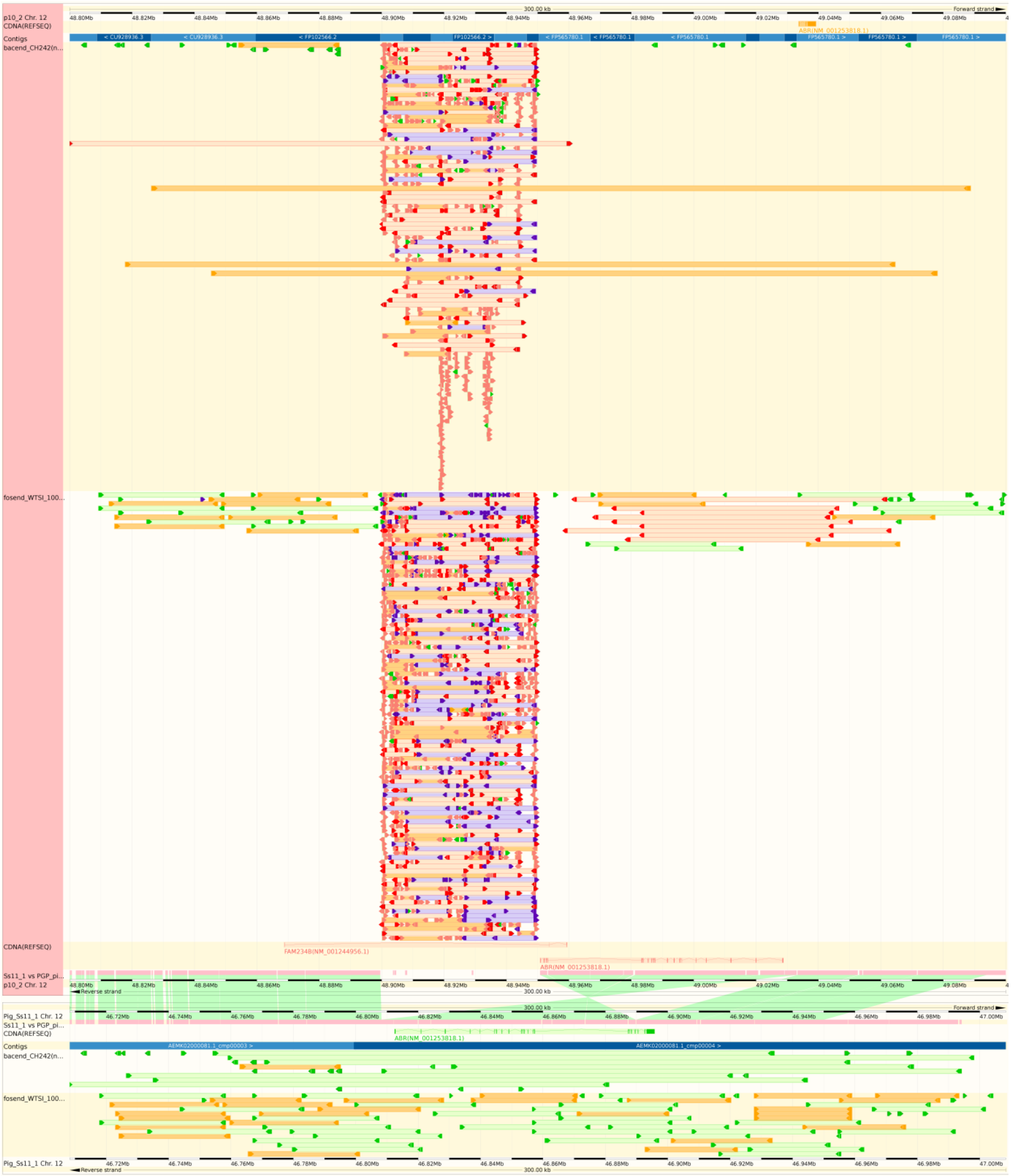

**Fig. S5. Assembly comparisons in gEVAL (SSC15).** Screenshot of gEVAL comparison of Sscrofa10.2, Sscrofa11.1 and USMARCv1.0 at *COL3A1*, *COL5A2* loci. In the new assembly (Sscrofa11.1, middle row marked with pink vertical block) an improved gene model for COL5A2 can be annotated; in the previous assembly (Sscrofa10.2, upper row) the order and orientation of sequence contigs within BAC clone CH242-40P12 (ENA: FP339585.2) are not resolved. There is good agreement between the Sscrofa11.1 (middle row) and the USMARCv1.0 (lower row) although the USMARCv1.0 assembly of SSC15 is inverted relative to Sscrofa11.1.

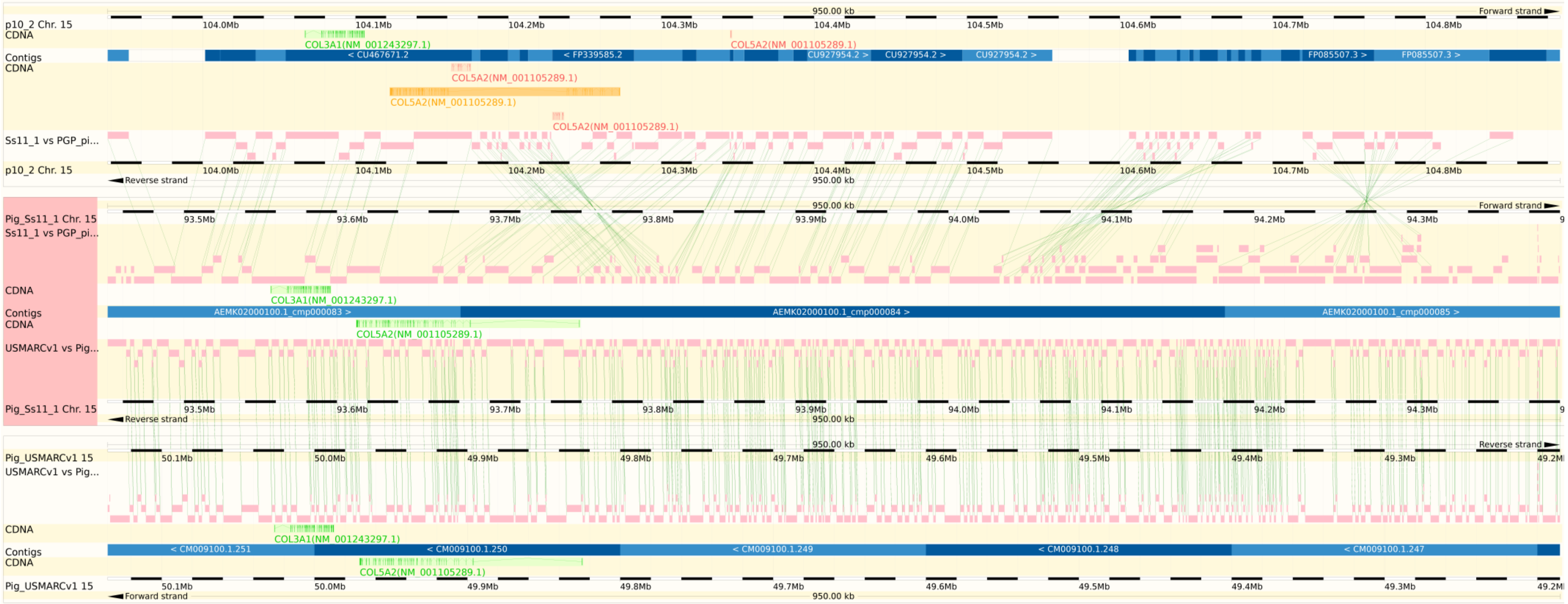

**Fig. S6. Assembly comparisons in gEVAL (SSC5).** Screenshot of gEVAL comparison of Sscrofa10.2, Sscrofa11.1 and USMARCv1.0 at the *KITLG* locus. The new assembly (Sscrofa11.1, middle row with pink vertical block at left hand side) resolves the sequences encoding *KITLG* which were split across two small scaffolds in Sscrofa10.2 (upper row). Although there is good agreement between Sscrofa11.1 (middle row) and USMARCv1.0 (lower row) assemblies in the right hand half of the region on SSC5 above, there is additional sequence present in the Sscrofa11.1 assembly between *DUSP6* and *KITLG*, the gene model for *KITLG* appears incomplete in the USMARCv1.0 assembly. Again the USMARCv1.0 is inverted relative to Sscrofa11.1.

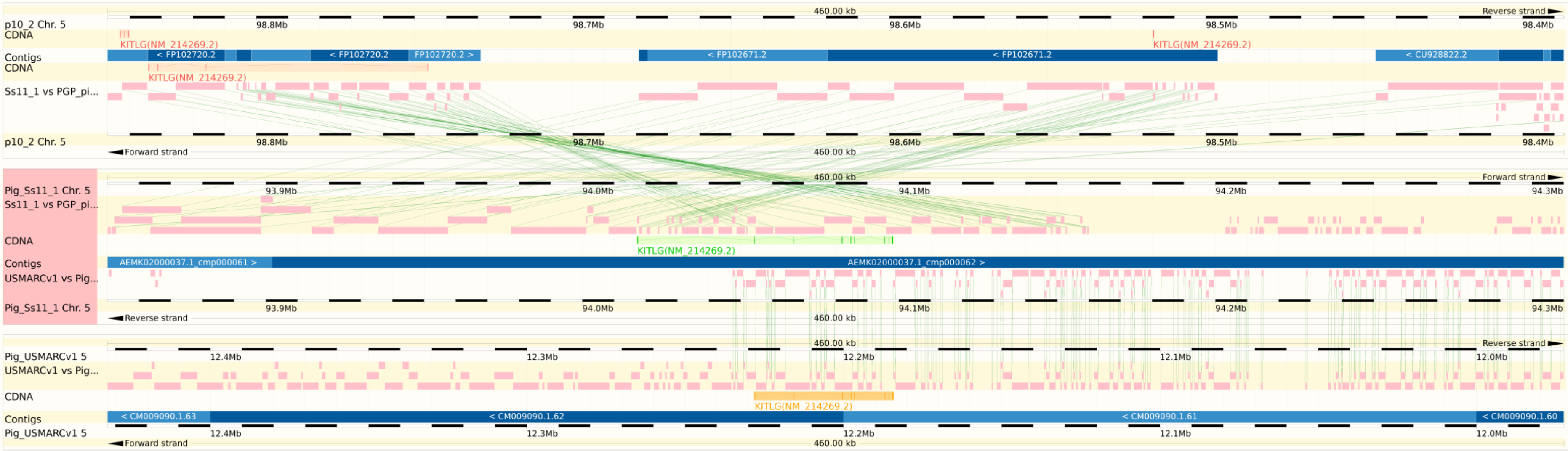

**Fig. S7. Assembly comparisons in gEVAL (SSC18).** Screenshot of gEVAL comparison of Sscrofa10.2, Sscrofa11.1 and USMARCv1.0 across the *ST7*, *CAPZA2* and *MET* loci on SSC18. The new assembly (Sscrofa11.1, middle row with pink block at left hand side) resolves the coding sequences for i) *ST7* that were previously split across two small scaffolds; *CAPZA2* that was similarly split across two small scaffolds; and iii) the *MET* sequences that were previously split as a result of an error in the orientation of the sequence drawn from BAC clone CH242-385N7 (ENA: CU633583.14) with respect to the sequence from BAC clone CH242-150K23 (ENA: CU694675.2) that harbours parts of the *MET* locus. This error in the incorporation of the CH242-385N7 (ENA: CU633583.14) in the Sscrofa10.2 assembly (upper row) is particularly unfortunate as this BAC had been sequenced to finish quality. There is good agreement between the Sscrofa11.1 (middle row) and USMARCv1.0 (lower row) assemblies with both SSC18 assemblies also being in the same orientation.

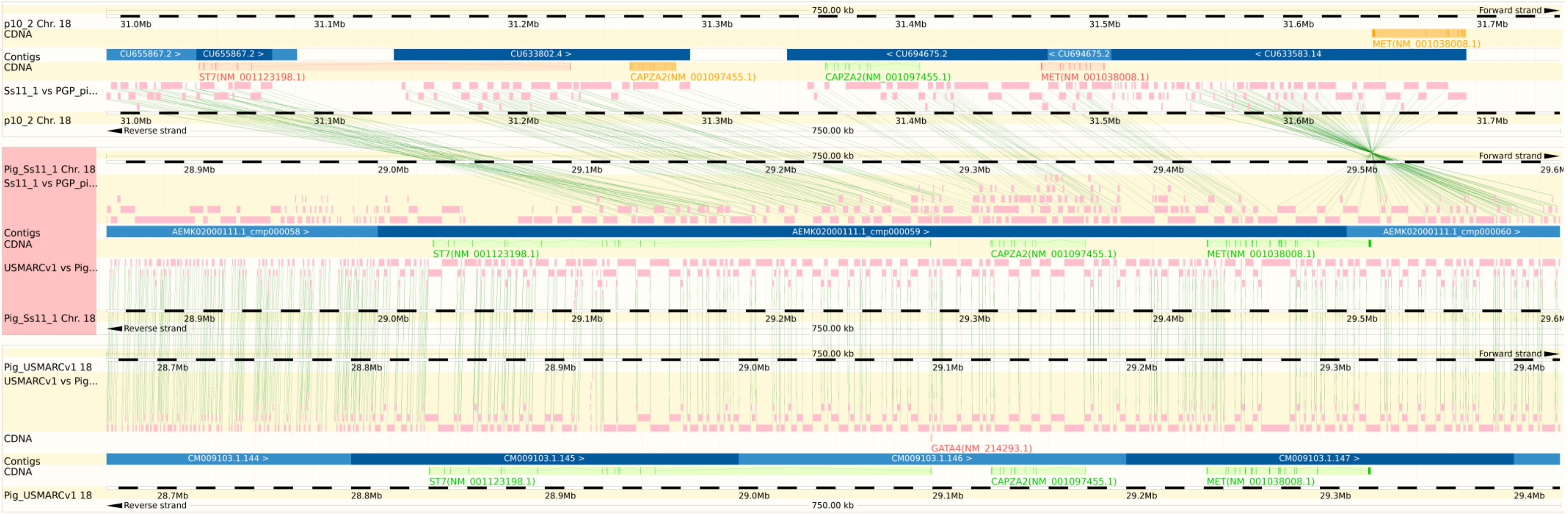

**Fig. S8. Order and orientation of SSC18 assemblies. A.** alignment of Sscrofa11.1 and USMARCv1.0 assemblies of SSC18; **B.** alignment of Sscrofa11.1 and radiation hybrid map (RH2); **C.** alignment of USMARCv1.0 and radiation hybrid map (RH2). The matches shown in the grey zone at the top of each plot of the Sscrofa11.1 versus USMARCv1.0 alignments probably represent a mix of repetitive sequences and matches to the unplaced scaffolds in the USMARCv1.0 assembly.

**A.** **B.** **C.**

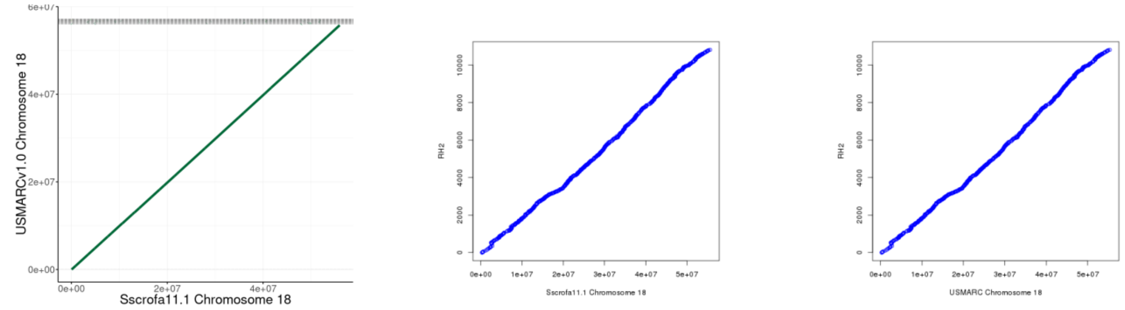

**Fig. S9**. **Order and orientation of SSC7 assemblies. A**. alignment of Sscrofa11.1 and USMARCv1.0 assemblies of SSC7; **B.** alignment of Sscrofa11.1 and radiation hybrid map (RH2); **C.** alignment of USMARCv1.0 and radiation hybrid map (RH2). The matches shown in the grey zone at the top of each plot of the Sscrofa11.1 versus USMARCv1.0 alignments probably represent a mix of repetitive sequences and matches to the unplaced scaffolds in the USMARCv1.0 assembly.

A. B. C.

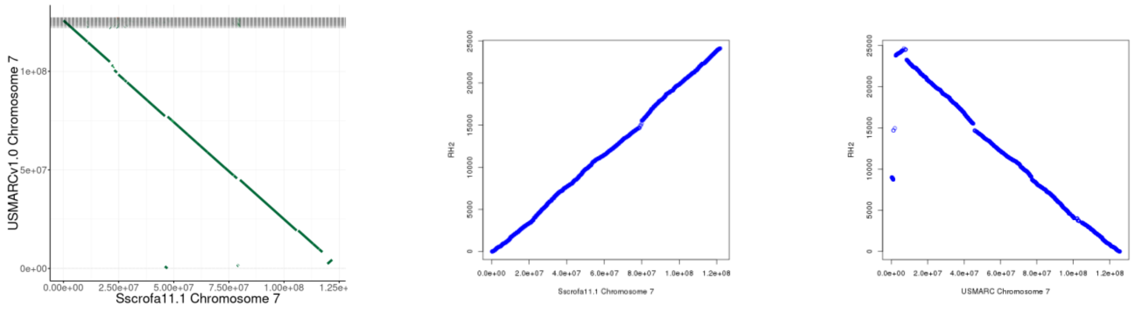

**Fig. S10. Order and orientation of SSC8 assemblies. A.** alignment of Sscrofa11.1 and USMARCv1.0 assemblies of SSC8; **B.** alignment of Sscrofa11.1 and radiation hybrid map (RH2); **C.** alignment of USMARCv1.0 and radiation hybrid map (RH2). The matches shown in the grey zone at the top of each plot of the Sscrofa11.1 versus USMARCv1.0 alignments probably represent a mix of repetitive sequences and matches to the unplaced scaffolds in the USMARCv1.0 assembly.

**A.** **B.** **C.**

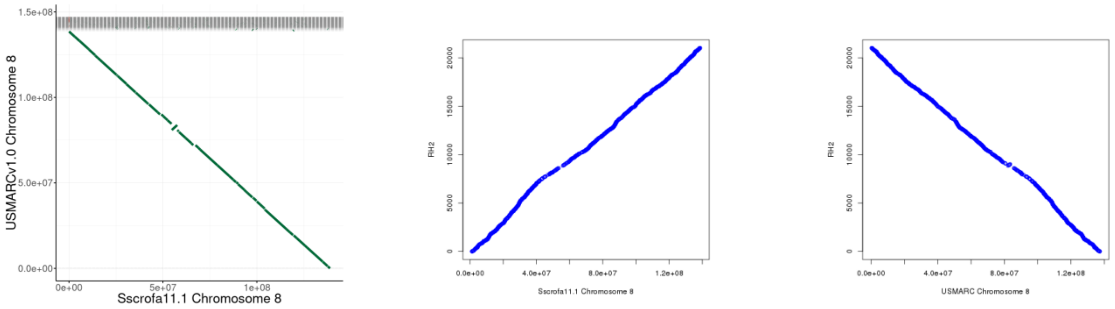

**Fig. S11. Assembly alignments.** Alignment of Sscrofa11.1 and USMARCv1.0 assemblies after correcting inversions of USMARCv1.0 chromosome scaffolds

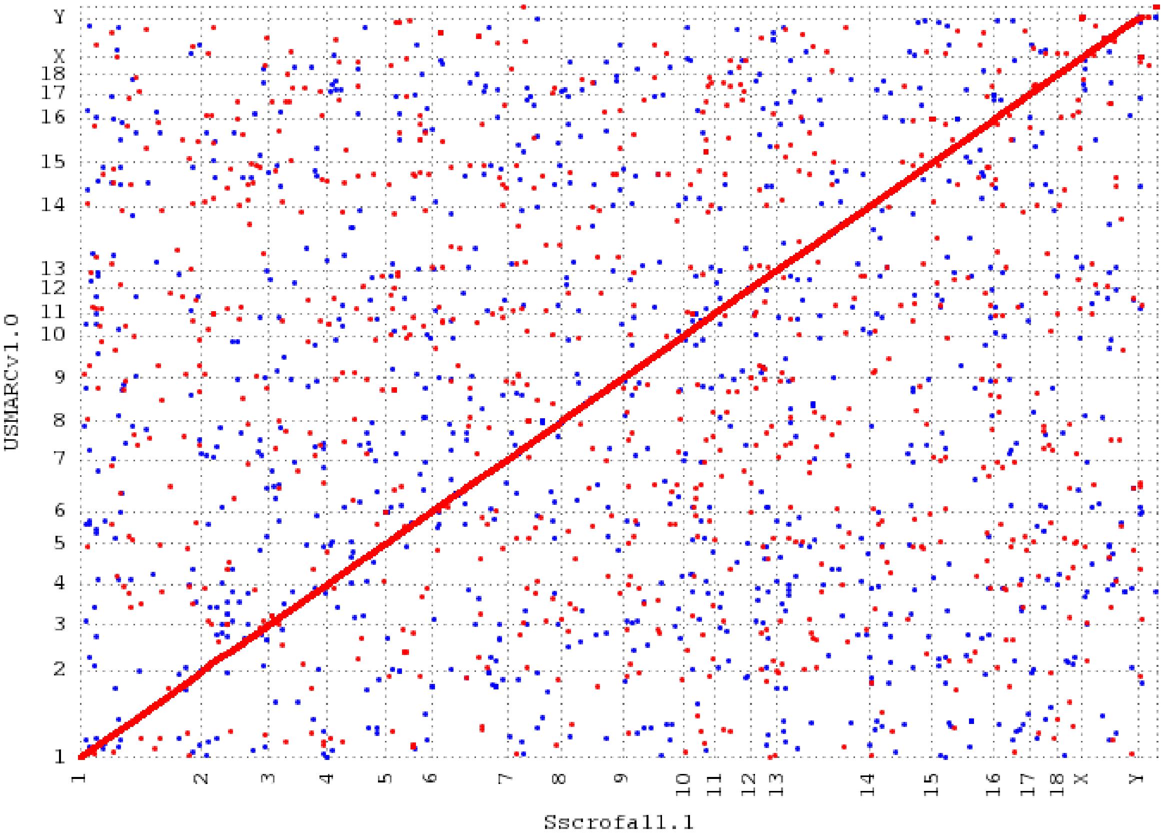

**Figure S12a. Assemblytics results.** Assemblytics comparison of Sscrofa11.1 (query) against the Sscrofa10.2 (reference) i). (left hand panel) variants from 50 to 500 bp; ii). (right hand panel) variants from 500 to 10,000 bp.

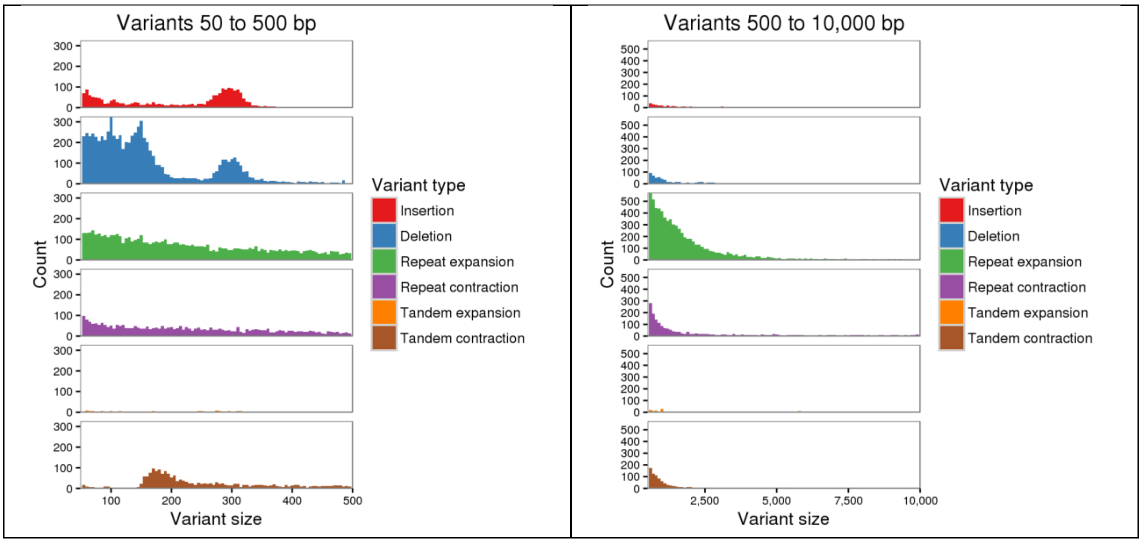

**Figure S12b. Assemblytics results.** Assemblytics comparison of USMARCv1.0 (query) against the Sscrofa10.2 (reference) i). (left hand panel) variants from 50 to 500 bp; ii). (right hand panel) variants from 500 to 10,000 bp.

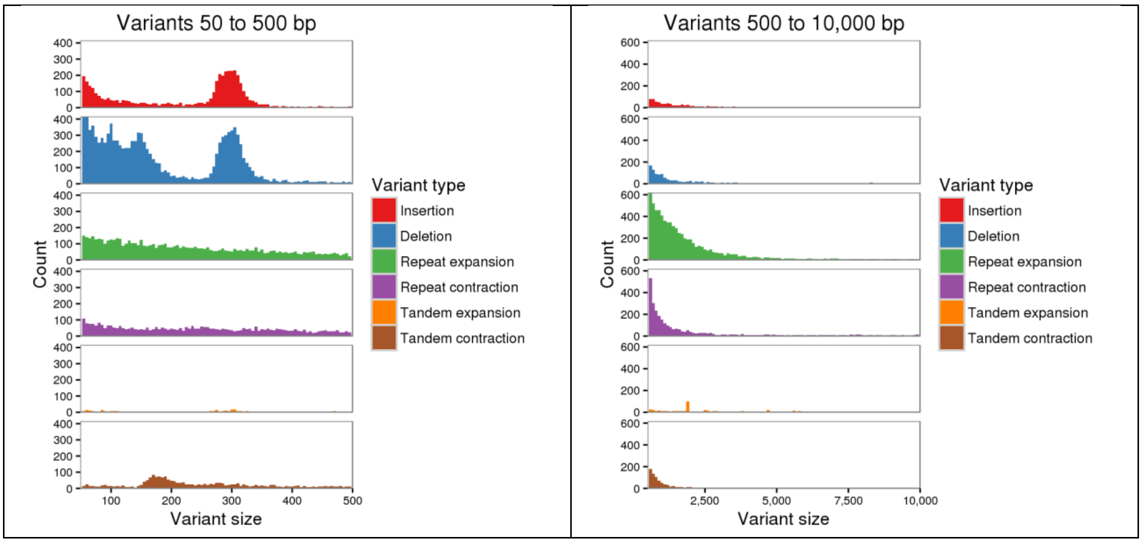

**Figure S12c. Assemblytics results.** Assemblytics comparison of USMARCv1.0 (query) against the Sscrofa11.1 (reference) i). (left hand panel) variants from 50 to 500 bp; ii). (right hand panel) variants from 500 to 10,000 bp.

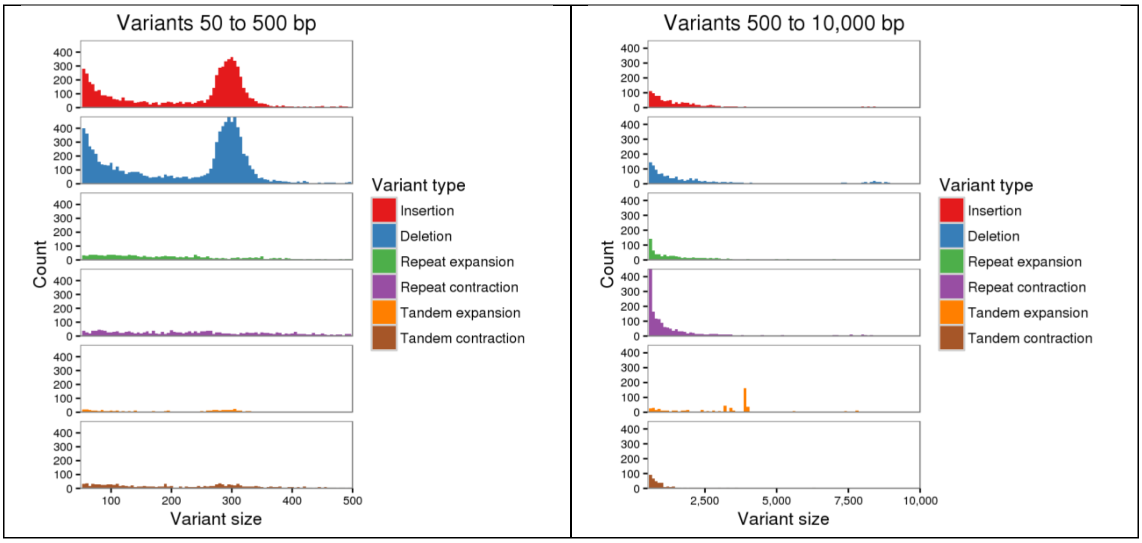

**Figure S13. Counts of repetitive elements in four pig assemblies.** Counts are given for repeat classes for which percent divergence was less than 40% and mapped length was above 70% relative to the RepBase database entries.

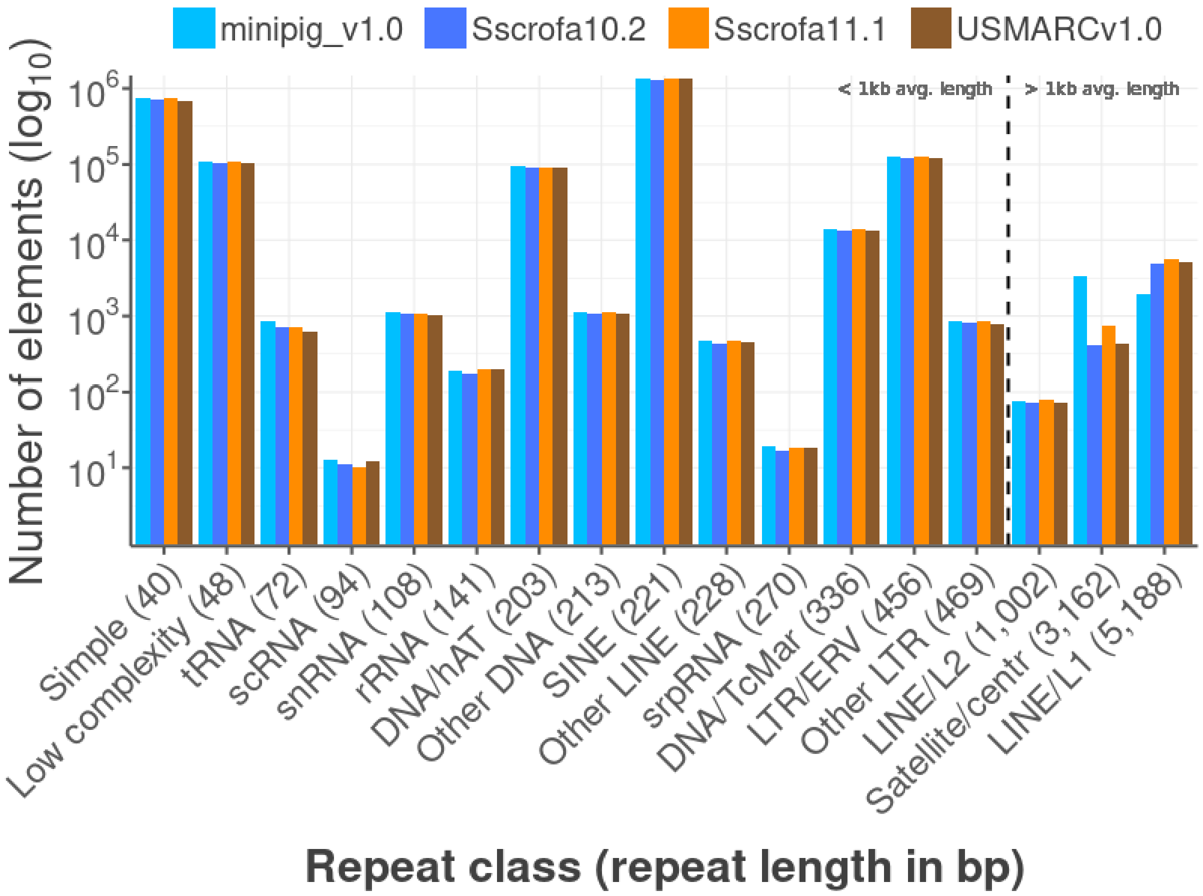

**Figure S14. Average mapped length of repetitive elements in four pig genomes.**

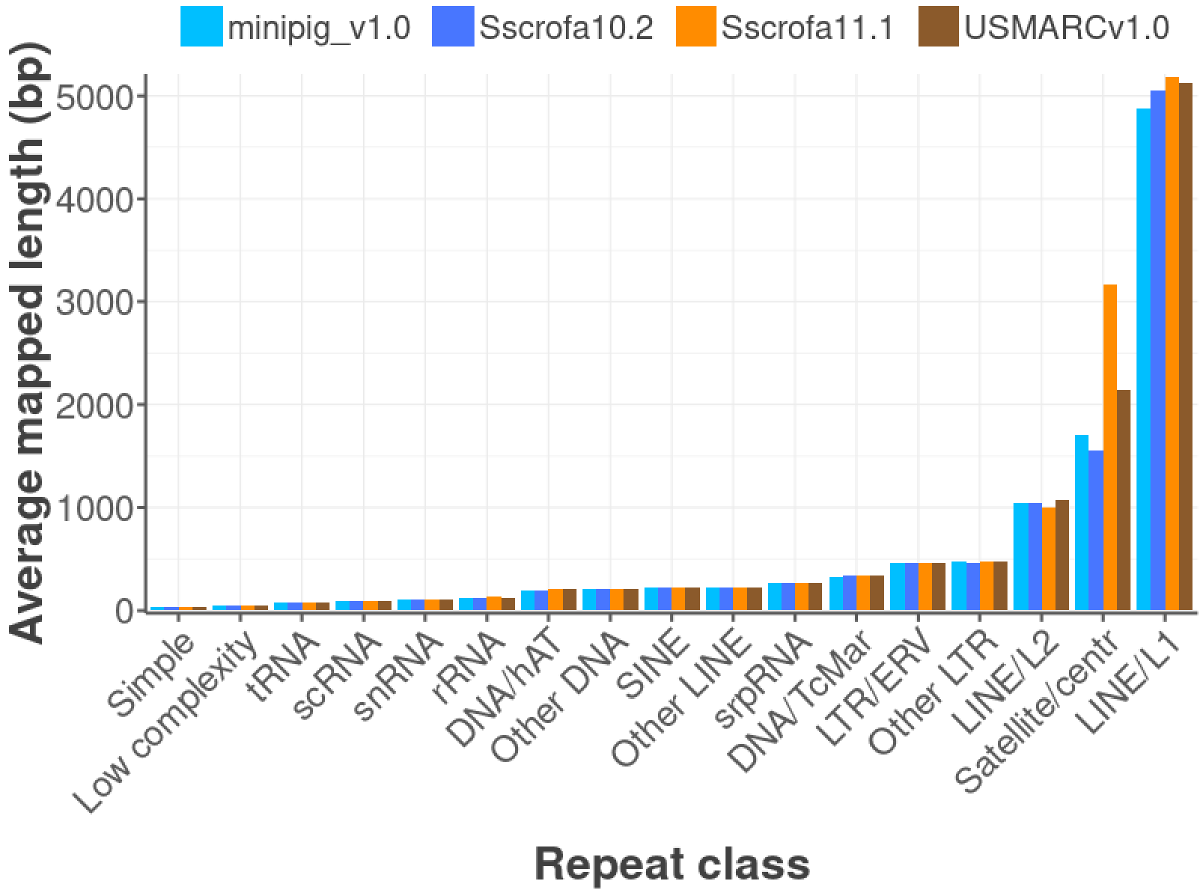

**Figure S15. Assembly SNP rank concordance versus reported chromosomal location.**

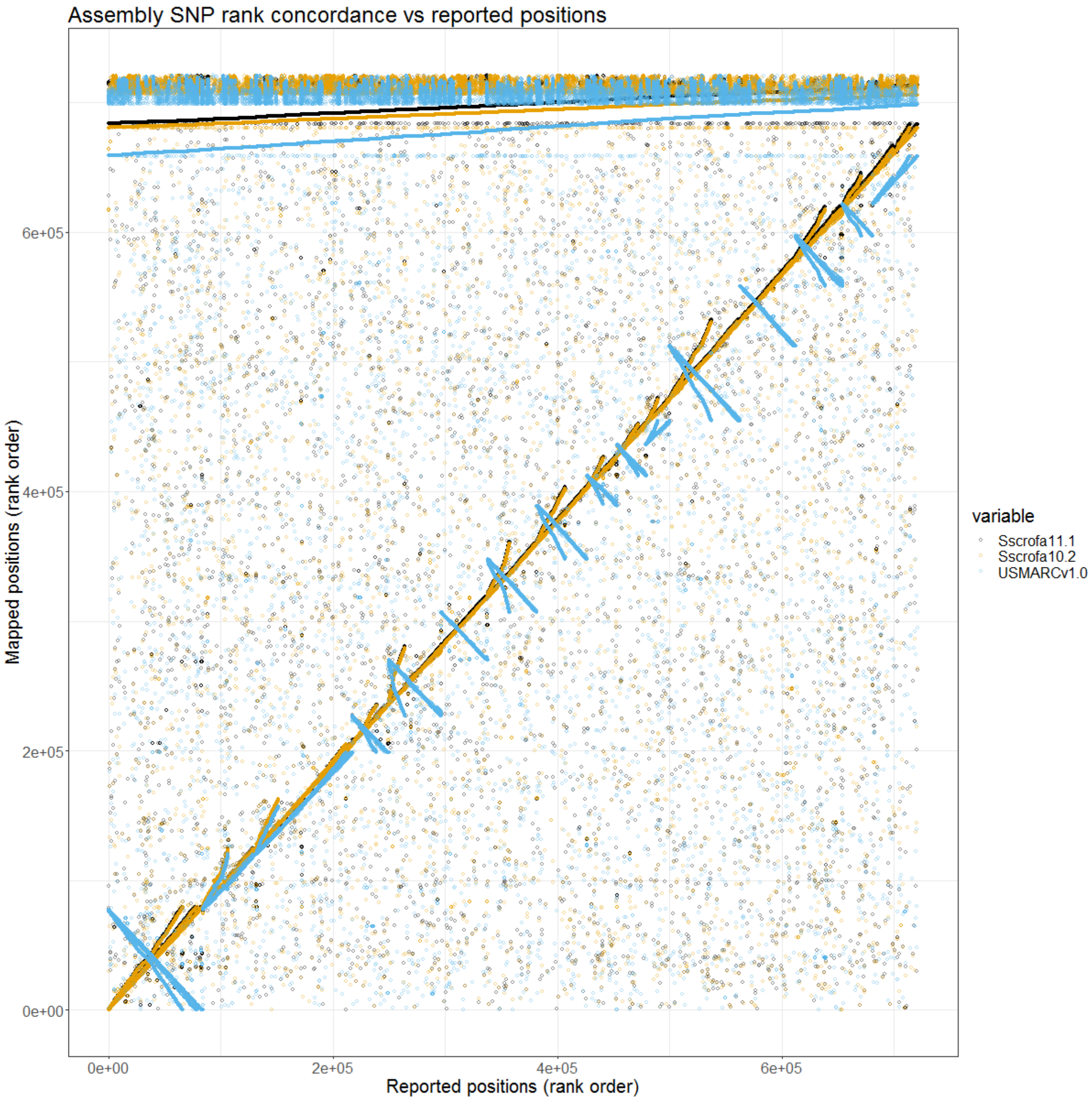
